# TAC-C uncovers open chromatin interaction in crops and SPL-mediated photosynthesis regulation

**DOI:** 10.1101/2025.02.10.637364

**Authors:** Jingmin Kang, Zhaoheng Zhang, Xuelei Lin, Fuyan Liu, Yali Song, Peng Zhao, Yujing Lin, Xumei Luo, Xiaoyi Li, Yanyan Li, Wenda Wang, Cuimin Liu, Shengbao Xu, Xin Liu, Jun Xiao

**Affiliations:** Institute of Genetics and Developmental Biology, Chinese Academy of Sciences, Beijing 100101, China; BGI Research, Beijing 102601, China; University of Chinese Academy of Sciences, Beijing 100049, China; College of Agronomy, Northwest A&F University, Yangling, Shaanxi 712100, China; Institute of Botany, Chinese Academy of Sciences, Beijing 100093, China; Centre of Excellence for Plant and Microbial Science (CEPAMS), JIC-CAS, Beijing 100101, China

## Abstract

*Cis*-regulatory elements (CREs) direct precise gene expression for development and environmental response, yet their spatial organization in crops is largely unknown. We introduce Transposase-Accessible Chromosome Conformation Capture (TAC-C), a method integrating ATAC-seq and Hi-C to capture fine-scale chromatin interactions in four major crops: rice, sorghum, maize, and wheat. TAC-C reveals that chromatin interaction frequency aligns with genome size and gene expression, exhibiting distinct loop structures between C3 and C4 plants, particularly in C4-specific enzymes coding genes. Integrating chromatin structure with population genetics data highlights that chromatin loops connect distal regulatory elements to phenotypic variation. Asymmetrical open chromatin interactions among subgenomes, driven by transposon insertions and sequence variations, contribute to biased homoeolog expression. Furthermore, TaSPL7/15 regulate photosynthesis-related genes through chromatin interactions, with enhanced photosynthetic efficiency and starch content in *Taspl7&15* mutant. TAC-C provides new insights into the spatial organization of regulatory elements in crops, especially for SPL-mediated photosynthesis regulation in wheat.

**Teaser:** TAC-C Reveals Chromatin Interactions and SPL-Mediated Photosynthesis Regulation in Crops.

## Introduction

*Cis*-regulatory elements (CREs) are crucial for fine-tuning gene expression, influencing phenotypic diversity, and accelerating crop improvement (*1–4*). In rice, a 5.3 kb silencer upstream of the spike development gene *FRIZZY PANICLE* (*FZP*) is bound by Brassinazole 1 (*OsBZR1*), suppressing *FZP* expression and influencing spike architecture and yield (*5*). In maize, a distal region over 40 kb upstream of *TEOSINTE BRANCHED1* (*TB1*) act in *cis* to alter *TB1* transcription, impacting inflorescence structure (*6*). In sorghum, binding by WRKY and ZF-D (zinc finger-DHHC) transcription factors (TFs) activates the aluminum tolerance gene *multidrug and toxic compound extrusion* (*SbMATE*), enhancing tolerance (*7*). These active CREs are typically located in accessible chromatin regions (ACRs) and can act as proximal or distal elements, influencing gene expression locally or over long distances (*8*). Techniques such as Assay for Transposase Accessible Chromatin sequencing (ATAC-seq), DNase-seq, and Formaldehyde-Assisted Isolation of Regulatory Elements (FAIRE-seq) have effectively identified ACRs across plant genomes, though they do not capture their spatial organization (*9–11*). The number of ACRs correlates with gene density, and in larger genomes, distal ACRs—often functioning as enhancers—are abundant (*12*). These distal elements play a key role in regulating tissue-specific gene expression, particularly in species with large genomes (*13*).

Recent advancements, including Chromosome conformation capture (3C) coupled with sequencing (Hi-C), Chromatin interaction analysis by paired-end tag sequencing (ChIA-PET), *in situ* Hi-C followed by chromatin immunoprecipitation (HiChIP), open chromatin enrichment and network Hi-C (OCEAN-C), and chromatin interaction analysis by ATAC (ChIATAC), have significantly enhanced our understanding of chromatin interactions and three dimensional (3D) genome structures (*14–18*). The dynamics of chromatin loops are crucial for regulating organ development and stress responses in crops (*19, 20*). The structure of chromatin provides a physical framework for gene regulation, by reducing the physical distance between *cis*-regulatory elements and target genes, chromatin organization provides a framework for efficient gene regulation, especially in large genome like maize and wheat, where nuclear metabolic efficiency is vital (*21, 22*).

Polyploidy, common in crops (*23*), often leads to chromatin structure distinct from those in diploid species, with inter-subgenome chromatin interactions forming a novel regulatory layer that supports coordinated gene expression (*24, 25*). While Hi-C provides a broad view of genome-wide interactions, it may overlook critical contacts between open chromatin and distal regulatory elements (*26*). Techniques like ChIA-PET and HiChIP offer more specificity, focusing on interactions mediated by DNA-binding proteins or histone marks (*14, 17*). OCEAN-C integrates Hi-C with FAIRE-seq but lacks the signal clarity and TFs binding footprint resolution of ATAC-seq (*27, 28*). ChIATAC employs Sodium dodecyl sulfate (SDS) treatment of nuclei, producing data distinct from ATAC-seq (*18*). There remains a need for methods that better capture the spatial organization of open chromatin to deepen our understanding of long-range transcriptional regulation.

Common wheat (*Triticum aestivum*, *T. aestivum*) is a widely cultivated allohexaploid crop with a complex genome structure due to its polyploidization, comprising three subgenomes (A, B, and D) and a total genome size of 16G (*29*). This makes wheat an ideal model for exploring the 3D spatial organization of large-genome species. Studies using Hi-C and HiChIP have revealed that interactions within subgenomes are more frequent than those between subgenomes (*30*), indicating a higher-order organization favoring subgenome-specific affinities. ATAC-seq has identified numerous CREs in wheat that play vital roles in gene expression and phenotypic traits (*4, 31, 32*). However, the mechanisms by which distal CREs, particularly their long-range interactions, regulate gene expression remain poorly understood. High-resolution open chromatin interaction maps generated using OCEAN-C for wheat and its tetraploid and diploid relatives reveal more frequent chromatin interactions within the D subgenome (*28*), suggesting that it may have evolved more rapidly and retained more epigenetic features from its diploid ancestors. Despite advancements, critical questions remain, such as how specific quantitative trait locus (QTLs) and expression quantitative trait locus (eQTLs) regulate gene expression through long-range interactions, the role of chromatin interactions in homoeolog gene expression bias, and the mechanisms driving chromatin loop formation in wheat.

The establishment and maintenance of nuclear 3D structures have long posed challenges in 3D genome research (*33, 34*). In animals, the chromatin loop formation is often mediated by CCCTC-binding factor (CTCF) and cohesin complexes (*35–37*). The TF Zinc Finger Protein 143 (ZNF143) in animals has been shown to work in tandem with CTCF, regulating CTCF/cohesin interactions and influencing topologically associating domains (TADs) formation (*38*). The lack of a CTCF homolog in plants suggests that any TAD-like structures observed in plants may have evolved through independent pathways (*39*). Recent studies have indicated that in rice (*39*), members of the TCP (Teosinte branched1/Cycloidea/Proliferating cell factor) family of TFs may play a role similar to CTCF in animals. TCPs have been found to recognize specific motifs at the boundaries of rice TAD-like structures (*39*). Additionally, TCP TF activity has been linked to 3D chromatin structure regulation in the liverwort *Marchantia* (*40*). Despite these advances, research into TFs mediating 3D genome structures in wheat remains limited. Investigating the role of specific wheat TFs in nuclear architecture could provide valuable insights into plant genome regulation and evolutionary parallels between plant and animal 3D genome structures.

In this study, we developed the Transposase-Accessible Chromosome Conformation Capture (TAC-C) technique, integrating *in situ* Hi-C and ATAC-seq to map fine-scale open chromatin interactions in wheat, rice, maize and sorghum. We explored the regulatory effects of distal open chromatin on gene expression, focusing on asymmetrical open chromatin interactions within different wheat subgenomes. Additionally, we uncovered the key role of SBP (SQUAMOSA promoter binding protein) family TFs in shaping chromatin interactions and their significant impact on regulating photosynthetic energy metabolism in wheat. These findings provide valuable insights and resources for studying 3D genome architecture and transcriptional regulation in complex crop genomes.

## Results

### TAC-C efficiently capture fine-scale chromatin interactions

To capture chromatin interactions at accessible regions and explore long-range transcriptional regulatory processes in large-genome species with low sequencing depths, we developed a new strategy. This method integrates ATAC-seq with Hi-C, resulting in a more efficient technique we term Transposase-Accessible Chromosome Conformation Capture (TAC-C). Plant tissue was first fixed with formaldehyde, followed by the extraction of high-quality nuclei. To preserve open chromatin regions, we avoid SDS treatment and instead perform *in situ* restriction enzyme digestion with DpnII, followed by biotin labeling and proximity ligation, slightly differing from traditional *in situ* Hi-C protocol. Next, we apply transposase for *in situ* tagmentation, as per the ATAC-seq protocol. Biotin-labeled chromatin ligation products are isolated using biotin-streptavidin affinity, and the TAC-C library is generated through PCR (**Fig. 1A**, see Methods for details).

**Fig. 1.**
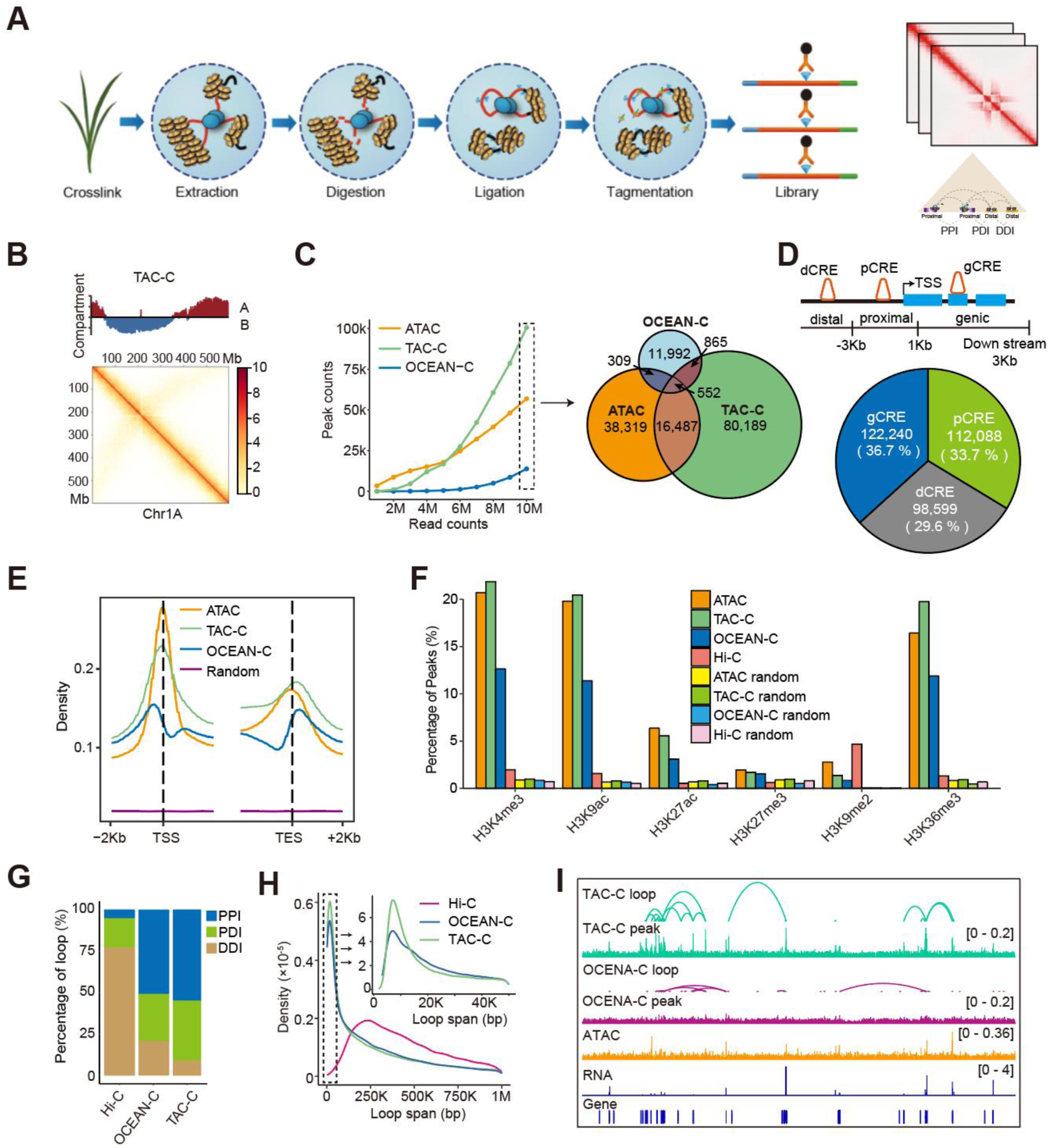
Identification of open chromatin interactions using TAC-C and comparison with other techniques. **A.** Schematic of TAC-C. **B.** TAC-C interaction matrix at 1Mb resolution across chromosome 1A, with chromatin compartments shown as a track above the contact map. Compartment A is marked in blue, and compartment B in red. **C.** Statistical comparison of peak capture efficiency under different sequencing depth using TAC-C, ATAC, and OCEAN-C. Venn diagram showing the number of peaks captured by each method based on 10M of sequencing reads. **D.** Classification of CREs based on their distribution around genes, with the number and proportion of each CRE type of CRE indicated. TSS: transcription start site, dCRE: distal CRE, pCRE: proximal CRE, gCRE: genic CRE. **E.** Density plot of TAC-C, OCEAN-C, ATAC, and random regions around transcription start site (TSS) and transcription termination site (TTS). **F.** Percentage overlap of TAC-C, OCEAN-C, Hi-C, and ATAC peaks or random regions with various histone modification marks. **G.** Percentage of proximal-proximal interactions (PPI), proximal-distal interactions (PDI), and distal-distal interactions (DDI) identified by TAC-C, OCEAN-C and Hi-C. **H.** Loop span distribution of open chromatin interactions identified by TAC-C, OCEAN-C and Hi-C. Regions with loop span below 50kb are highlighted in the top right corner. **I.** Example of open chromatin interactions: browser view of a 6.6Mb region showing a randomly selected open chromatin interactions with associated TAC-C and OCEAN-C data.

The TAC-C data exhibited high reproducibility between biological replicates (**fig. S1A, table S1**). The TAC-C interaction map at the chromosome level exhibited robust signals along the primary diagonals and anti-diagonal lines (**Fig. 1B and fig. S1B**). This pattern reflected the Rabl-like chromosome configuration, consistent with the Hi-C interaction matrices as reported (*30*), indicating that TAC-C data can effectively depict the high-order organization of the wheat genome. We then evaluated the effect of sequencing depth on the TAC-C peak calling and compared it against OCEAN-C and ATAC-seq techniques (*16, 41*). As the read count increased, TAC-C generated peaks similar to or even more than ATAC-seq, largely outperforming OCEAN-C. Specifically, ∼30% of ATAC-seq peaks overlapped with TAC-C, while less than 2% overlapped with OCEAN-C at 10 million reads sequencing depth (**Fig. 1C**), suggesting that TAC-C more effectively capture open chromatin regions.

In wheat, we identified 332,927 overlapping TAC-C peaks from two biological replicate libraries (Rep1: 398 million reads; Rep2: 405 million reads) for subsequent analysis. These peaks were categorized as gene body (g, 36.7%), promoter (p, 33.7%), and distal (d, 29.6%) CREs (**Fig. 1D**). TAC-C and ATAC-seq peaks were strongly enriched at transcription start sites (TSS), largely more than OCEAN-C peaks (**Fig. 1E**). Correlation analysis with epigenetic markers revealed that TAC-C peaks are predominantly associated with active histone modifications (H3K4me3, H3K9ac, and H3K27ac), similar to ATAC-seq and remarkably higher than OCEAN-C and Hi-C peaks (*30, 41*) (**Fig. 1F**). This indicates that TAC-C peaks are mostly located in active *cis*-acting elements, such as promoters or enhancers. We further compared TAC-C with OCEAN-C and Hi-C in identifying open chromatin interactions. TAC-C identified 159,667 intrachromosomal loops (**table S2**), classified into proximal anchors (P) within 3 kb distance to TSS of genes and distal anchors (D) beyond 3 kb. The loops consisted of 55.5% P-P interactions (PPIs), 35.5% P-D interactions (PDIs), and 9.0% D-D interactions (DDIs) (**Fig. 1G**). TAC-C showed a higher proportion of P-P interaction and detected shorter loop spans compared to OCEAN-C and Hi-C (**Fig. 1H**), indicating TAC-C’s superior performance in detecting fine-scale open chromatin interactions (**Fig. 1I**).

In summary, we have developed the TAC-C assay by integrating *in situ* Hi-C and ATAC-seq protocols, validating it experimentally in bread wheat. This method enhances the ability to capture open chromatin interactions, particularly in large-genome crops like wheat.

### Conserved and distinct features of chromatin interactions across four crop species

To explore the characteristics of interacted open chromatins in crops, we also conducted TAC-C experiments on young leaves from Rice (*Oryza. Sativa*, *O. sativa*), Sorghum (*Sorghum. Bicolor*, *S. bicolor*) and Maize (*Zea. Mays*, *Z. mays*), observing high reproducibility between biological replicates (Pearson correlation 0.91∼0.99) (**fig. S1A**). The number of identified loops ranged from 46,327 to 159,667 across species, correlating well with the number of annotated genes within each species (**Fig. 2A and fig. S2A**). Larger genomes (*Z. mays* and *T. aestivum*) displayed notably longer distances for open chromatin interactions compared to smaller genomes (*O. sativa* and *S. bicolor*), with a greater percentage of PDI and DDI (**Fig. 2B and fig. S2B**), where the percentage of PDI closely correlated with genome size (**fig. S2C**).

**Fig. 2.**
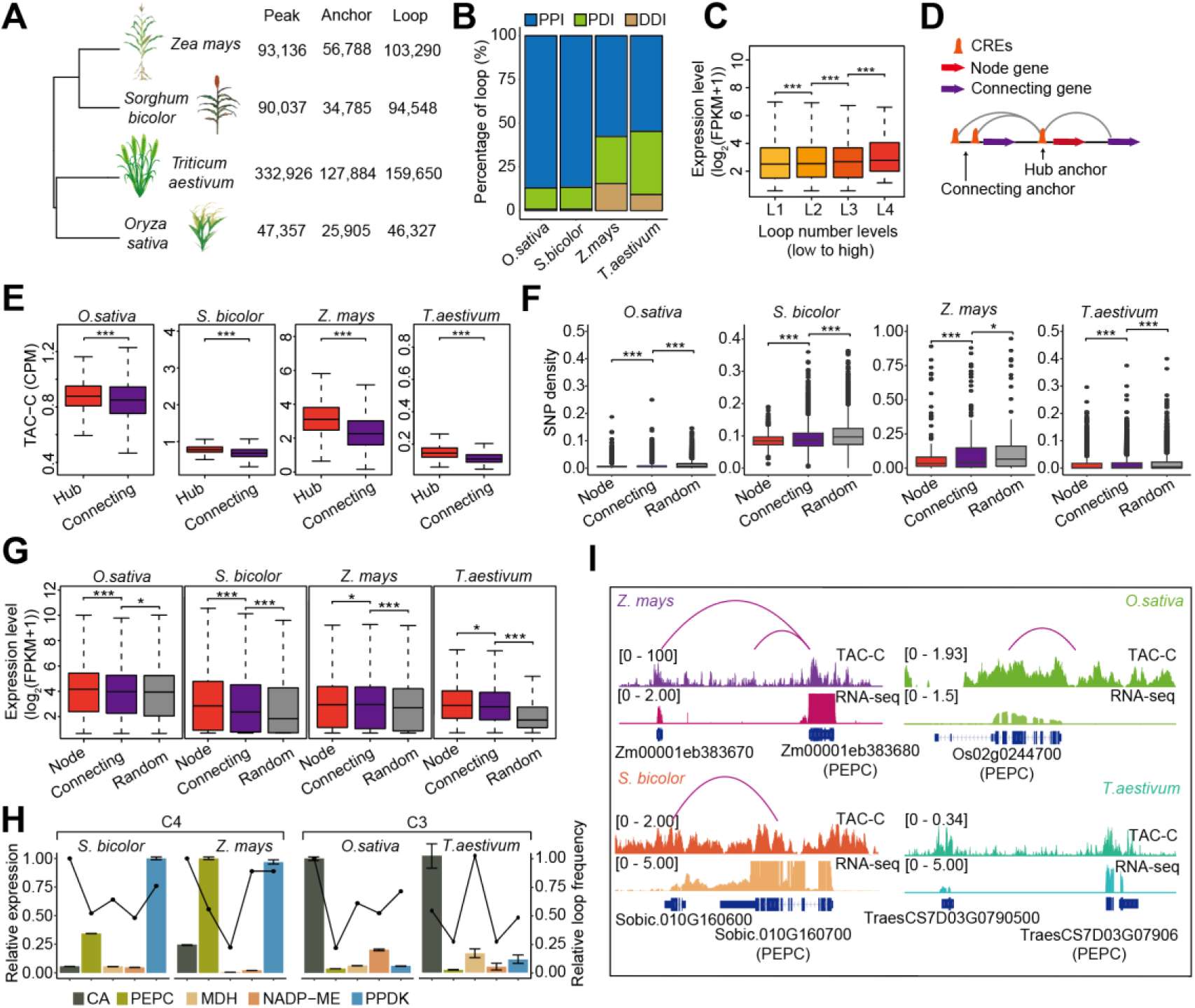
Characteristics of chromatin interactions across four crops species. **A.** Phylogenetic tree of four crop species with the number of peaks, anchors, and loops identified from TAC-C data. **B.** Percentage distribution of PPI, PDI, and DDI across the four crop species. **C.** Expression levels (log_2_(FPKM+1)) of genes with varying levels of chromatin interactions in *T. aestivum*. ***, *p*<0.001 from Student’s *t*-test. **D.** Schematic representation of the hub-and-connection anchor model of chromatin interactions, with definitions of for node and connecting genes based on contact frequency. **E.** TAC-C signal intensity in hub and connecting anchor regions across the four crop species. ***, *p*<0.001 from Student’s *t*-test. **F-G.** SNP density (**F**) and expression levels (**G**) in node, connecting and random genes across the four crop species. *, *p*<0.05, **, *p*<0.01, ***, *p*<0.001 from Student’s *t*-test. **H.** Relative expression levels and loop frequency in key C4 pathway gene (CA, PEPC, MDH, NADP-ME, PPDK) in C4 (*S. bicolor* and *Z. mays*) and C3 (*O. sativa* and *T. aestivum*) plants. **I.** Browser view showing differences in open chromatin interaction and expression levels for *PHPC* across the four crop species.

Chromatin loops have been implicated in gene expression regulation (*42, 43*). We found that gene pairs looped together tended to exhibit high expression levels simultaneously (**fig. S2D**). When genes were categorized based on chromatin interaction frequencies, those with more interactions showed higher expression levels across all four species (**Fig. 2C and fig. S2E**), suggesting a positive correlation between chromatin interaction frequencies and gene expression. The top 10% most frequently interacting anchors were defined as “hub anchors”, their partners as “connecting anchors”, whereas, other open chromatin peaks that were not located in the loop anchor were defined as basal peaks. Hub anchor-linked genes termed as “node genes” and connecting anchor-linked genes as “connecting genes” (*44*) (**Fig. 2D and fig. S2F**). Hub anchors exhibited higher TAC-C signals (**Fig. 2E**), while node genes shown lower SNP densities and higher expression levels across all four species (*45–48*) (**Fig. 2F and 2G**), underscoring the conservation and importance of hub anchors and node genes.

We further categorized the four species into C4 plants (*S. bicolor* and *Z. mays*) and C3 plants (*O. sativa* and *T. aestivum*), identifying 12,131 and 4,473 inter-species conserved homologous gene-pair, respectively, with 397 loops common to all four species (**fig. S2G**). Gene Ontology (GO) enrichment analysis revealed that pathways related to carbohydrate and energy metabolism are conserved within the homologous open chromatin interaction gene pairs in both C4 and C3 plants (**fig. S2H**). Five C4-specific enzymes, [CA (carbonic anhydrase, NCBI: EU970742), PEPC (phosphoenolpyruvate carboxylase 1, NCBI: NM_001111948), MDH (malate dehydrogenase, NCBI: EU964066), NADP-ME (NADP-dependent malic enzyme, NCBI: EU975279), and PPDK (phosphate dikinase 1, NCBI: NM_001112268)], play crucial roles in C4 carbon assimilation (*49*). The expression levels and loop frequencies of these genes showed significant difference between C4 and C3 plants, particularly *PEPC* and *PPDK*, which had higher expression and loop frequencies in C4 plants (**Fig. 2H and table S3**), highlighting the role of open chromatin interactions in the C4 pathway. For instance, *PEPC* was looped with its neighboring gene in both *S. bicolor* and *Z. mays*, but this loop was absent in *O. sativa* and *T. aestivum* (**Fig. 2I**).

Taken together, the length and number of open chromatin interactions correlate with genome size and gene count across species, with interaction frequency significantly influencing gene expression. Hub anchors display higher TAC-C signals, while node genes exhibit elevated expression levels and conserved sequences, indicating their significant biological roles. Notably, loop structures within C4 pathway-specific enzymes differ markedly between C3 and C4 plants.

### Chromatin interaction link regulatory elements to phenotype variation in wheat

To explore the importance of spatial organization of chromatin, we compiled data from published sources on QTLs and eQTLs related to various agronomic traits (**table S4**), as well as breeding selection sweeps (SS) in wheat to investigate whether chromatin interactions could spatially link distal regulatory elements to target genes, ultimately contributing to phenotypic variation (*50–53*). Our analysis revealed that hub anchors, which primarily localize to the distal ends of chromosomes, showed a distribution pattern consistent with “functional regions” as defined by QTLs, eQTLs and SS (**Fig. 3A**). Notably, hub anchors exhibited the highest overlap with these functional regions, followed by connecting anchors and basal peaks (**Fig. 3B**), suggesting that regions frequently involved in chromatin loops may play more crucial roles in gene regulation. Furthermore, QTLs overlapped with the hub anchor exhibit stronger significance, while eQTLs and genes looped by the hub anchor demonstrate larger effect sizes (*51, 54*) (**Fig. 3C and 3D**). This indicates a more pronounced role of regulatory elements within the hub anchor.

**Fig. 3.**
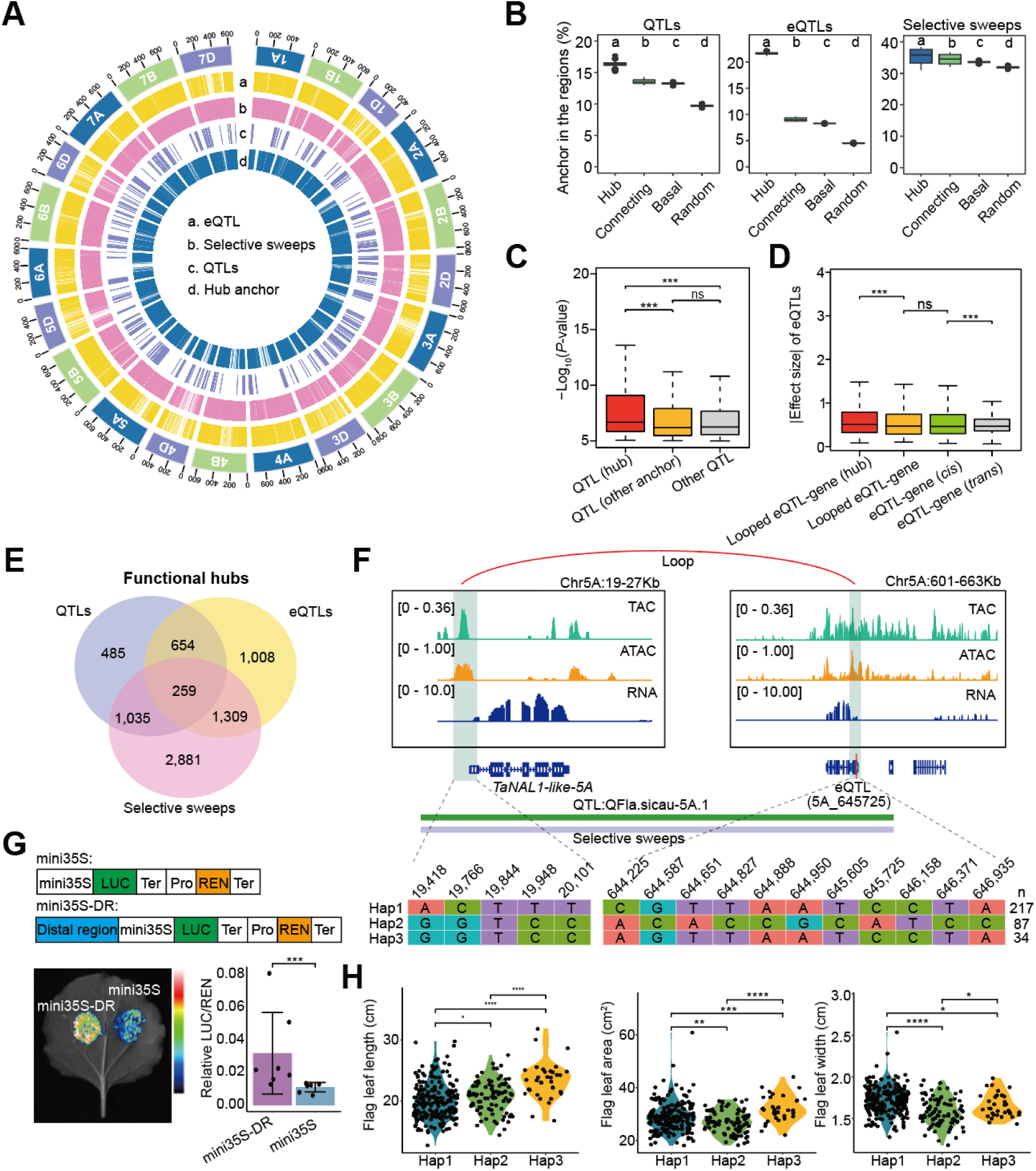
Long-range chromatin interactions regulate gene expression and associate with trait variations. **A.** Circos plot showing the distribution of wheat eQTLs (a), breeding selective sweeps (b), QTLs (c), and hub anchors (d) across the wheat genome. **B.** Percentage of overlapping interactions between hub anchors, connecting basal anchors, and random regions with QTLs, eQTLs and breeding selective sweep. To reduce false positives, 1000 permutation tests were conducted. In each test, 60% of accessions were randomly sampled, and differences were calculated using least significant difference (LSD) test. **C.** Significance of GWAS signals for SNPs located in hub anchor, other anchors and QTL regions. GWAS signals data were obtained from published sources (details in table S4). ***, *p*<0.001, ns, *p*>0.05 from Student’s *t*-test. **D.** Absolute effect size of looped eQTL-gene pairs compared to other eQTL-gene pair. Looped eQTL-gene (hub) pairs refer to those where the eQTL is located in the anchors or hub anchors. Effect size data were sourced from published studies (details in table S4). ***, *p*<0.001, ns, *p*>0.05 from Student’s *t*-test. **E.** Venn diagram showing the overlap between hub anchors, QTLs, eQTLs, and breeding selective sweeps. **F.** Open chromatin loop between *TaNAL1-like-5A* and distal eQTL (5A_645725). Green and pink tracks indicate QTL and breeding selection sweeps regions. The table below shows haplotypes of SNP sites in *TaNAL1-like-5A* and the distal regulatory region. **G.** Luciferase reporter assays validating the transcriptional regulatory role of the distal region on *TaNAL1-like-5A*. The distal anchor sequences, containing the eQTL locus, was introduced into the proximal region of *TaNAL1-like-5A* promoter (Distal region). ***, *p*≤0.001 from Student’s *t*-test. **H.** Comparison of flag leaf length, width and leaf area of *TaNAL1-like-5A* in three haplotypes of the loop-linked *TaNAL1-like-5A* and eQTL (5A_645725). *, *p*≤0.05; **, *p*≤0.01; ***, *p*≤0.001; ****, *p*≤0.0001 from Student’s *t*-test.

In total, 7,632 hub anchors overlapped QTLs, eQTLs and breeding selection sweeps. Of these, only 259 hub anchors were common across all three regions (**Fig. 3E**). For instance, an eQTL (Chr5A_645725) located 621 kb downstream of the leaf width-regulated gene *NARROW LEAF1* (*TaNAL1-like-5A*) was found within a flag leaf area (FLA)-regulated QTL and a breeding selection region (*51, 55, 56*). This eQTL, located in a hub anchor, was observed to loop with *TaNAL1-like-5A* (**Fig. 3F**). Furthermore, the functional potential of these distal regulatory regions was established through a luciferase reporter assay. *TaNAL1-like-5A* carrying the distal regions exhibited robust activity in the reporter assay, with higher signals compared with the controls carrying the 35S promoter alone (**Fig. 3G**). These findings suggest that distal regulatory elements can modulate target gene expression through chromatin interactions, thereby influencing phenotypic variation. To explore the genetic variation underlying this regulation, haplotype analysis was conducted for both *TaNAL1-like-5A* and the distal eQTL region Chr5A_645725 (*57*) (**Fig. 3F**). This analysis identified three distinct haplotypes, each exhibiting significant differences in flag leaf length, width and flag leaf area across various natural populations (*57*) (**Fig. 3H**).

Our findings reveal that distal elements may regulate the expression of target genes through chromatin interactions, thereby influencing phenotypic variation. Providing physical topological information support for distal regulatory sites identified through Genome-Wide Association Studies (GWAS) and Transcriptome-Wide Association Studies (TWAS).

### Asymmetric chromatin interaction associates with homoeolog expression bias in wheat

Since wheat is a hexaploid species, with proportion of A, B, and D subgenomes showing imbalanced expression pattern (*58*), we wonder whether differences in chromatin interactions among subgenomes contribute to this expression bias. We compared the loops at subgenomic levels for each chromosome. Interestingly, while the strength of the loops was comparable across three subgenomes, the D subgenome exhibited significantly more loops than the A and B subgenomes, consistent with previous study (*28*) (**fig. S3A, B**). In total, 39,714, 36,643, and 41,559 PPIs were identified in A, B, and D subgenomes, respectively. Among these, 3,742, 5,337 and 5,257 PPIs were conserved between two AB, AD and BD subgenomes, respectively. And 7,504 PPIs were conserved across all three subgenomes (**Fig. 4A**), indicating asymmetrical interactions between subgenome.

**Fig. 4.**
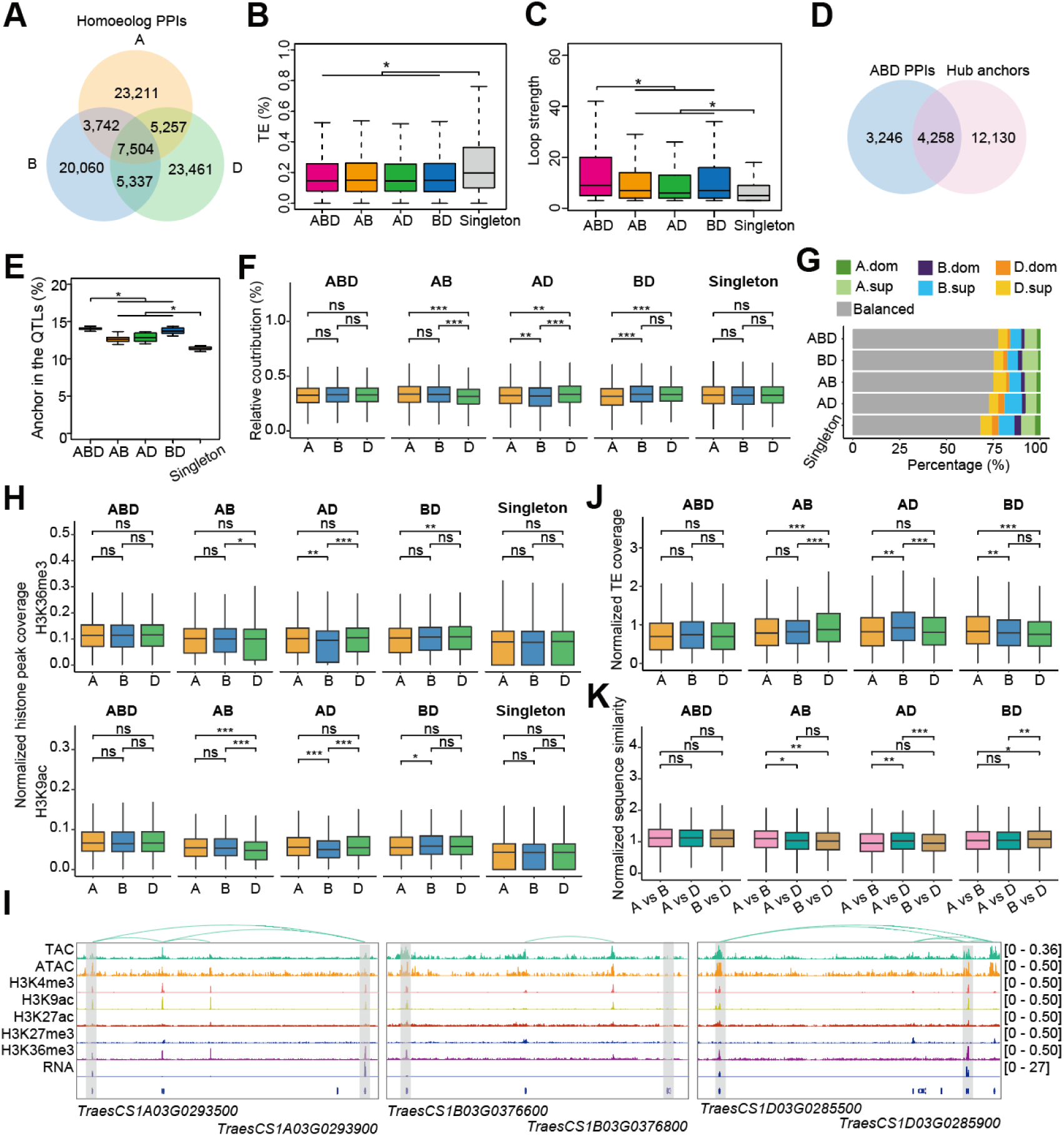
Asymmetrical open chromatin loops associated with homoeolog expression bias, epigenetic profiling, TE insertions, and sequence variations. **A.** Venn diagram showing the overlap of homoeolog PPIs between the A, B, and D subgenomes. **B-C.** Boxplots illustrating the TE coverage (**B**) and loop strength (**C**) across loop anchors at different homoeolog levels. ABD: homoeolog PPIs in all three subgenomes; AB, AD or BD: homoeolog PPIs in AB, AD or BD subgenomes; Singleton: PPIs in only one of the subgenomes A, B or D. *, *p*≤0.05 from Student’s *t*-test. **D.** Venn diagram showing the overlap between homoeolog ABD PPIs and hub anchors. **E.** Overlap ratio of QTL regions with PPI anchors at different homoeolog levels, normalized by the average QTL coverage in A, B and D subgenomes, respectively. **F.** Relative expression contribution of each subgenome based on triad, classified into five groups according to homoeolog PPI levels. *, *p*≤0.05; ***, *p*≤0.001 from Student’s *t*-test. **G.** Proportion of triads in each homoeolog expression bias category across the five homoeolog PPI groups. **H.** Coverage of H3K36me3 (upper) and H3K9ac (bottom) peaks in gene flanking regions (±5kb) across subgenomes, normalized to subgenome averages, classified by homoeolog PPI levels. *, *p*≤0.05; **, *p*≤0.01; ***, *p*≤0.001 from Student’s *t*-test. **I.** Browser screenshot showing asymmetrical open chromatin interactions and expression of linked gene (*TraesCS1A03G0293900*, *TraesCS1B03G0376800* and *TraesCS1D03G0285900*). Data tracks include TAC-C peaks, loops, histone modifications, ATAC profiling, and gene transcription. *, *p*≤0.05; **, *p*≤0.01; ***, *p*≤0.001 from Student’s *t*-test. **J.** TE coverage in gene flanking regions (±5kb) across subgenomes, normalized by the average TE coverage of singleton PPIs in A, B and D subgenomes. **K.** Pairwise comparisons of sequence similarity in gene flanking regions (±5kb) across subgenomes, normalized by the average gene sequence similarity between A vs B, B vs D, and A vs D, classified by homoeolog PPI levels. *, *p*≤0.05; **, *p*≤0.01; ***, *p*≤0.001 from Student’s *t*-test.

The coverage of transposable elements (TEs) in singleton loops was significantly higher than in homoeolog PPIs across subgenomes, suggesting a link between transposon insertion and the occurrence of asymmetrical interactions (**Fig. 4B**). The loop strength for ABD-conserved PPIs was the highest, while singleton PPIs exhibited the weakest loop strength (**Fig. 4C**). Additionally, 56.7% of loop anchors for ABD-conserved PPIs were classified as hub anchors, indicating that ABD-homoeolog interactions may represent crucial regulatory regions (**Fig. 4D**). Furthermore, homoeolog PPIs demonstrated a higher overlap with functional regions (**Fig. 4E**), and GO enrichment analysis revealed that genes associated with conserved PPIs are primarily involved in carbohydrate metabolism, photosynthesis, and growth and development (**fig. S3C**). In contrast, genes located in singleton loops were mainly enriched in basic cellular processes (**fig. S3D**).

To assess whether asymmetrical PPIs influence homoeolog expression bias, we compared the expression level of 16,539 triads containing at least one homoeolog linked with a loop. As expected, there were no significant differences in expression for ABD-conserved PPI-linked homoeologs. However, for PPIs conserved in only two subgenomes (AB, AD and BD), the expression levels of homoeolog lacking loops were significantly lower than the other two homoeologs (**Fig. 4F**). Furthermore, the proportion of balanced expression triads was highest in ABD-conserved PPI-linked genes. Homoeolog lacking loops were more likely to be suppressed in AB-, AD- or BD-conserved PPIs (**Fig. 4G**), indicating that asymmetrical interactions may be an important regulatory factor influencing homoeolog expression bias.

To explore the relationship between epigenetic modifications and asymmetrical PPIs, we analyzed histone modifications in the 5 kb regions flanking the asymmetrical loop-linked homoeolog triads. Activation histone marks, specifically H3K36me3 and H3K9ac, were correlated with the levels of PPI asymmetry. Homoeolog lacking loops exhibited lower coverage of these histone marks (**Fig. 4H**). Conversely, other epigenetic signals (ATAC, H3K27ac, H3K4me3, and H3K27me3) did not show significant difference (**fig. S3E**). For instance, a homoeolog triad (*TraesCS1A03G0293900*, *TraesCS1B03G0376800* and *TraesCS1D03G0285900*) looped with another homoeolog triad (*TraesCS1A03G0293500*, *TraesCS1B03G0376600* and *TraesCS1D03G0285500*) in A and D subgenomes, but this interaction was absent in the B subgenome (**Fig. 4I**). These asymmetrical loops were associated with elevated expression levels and activating chromatin modification signals in the A and D subgenomic homoeologs.

To further understand the factors driving asymmetrical PPIs, we compared TE coverage and sequence similarity in the 5 kb regions flanking the asymmetrical loop-linked homologous triads (**Fig. 4J and 4K**). The results showed that homoeolog lacking loops exhibited higher TE coverage and lower sequence similarity in their flanking regions. This suggests that transposon insertion or sequence variation may disrupt loop structure, leading to the loss of homologous loops in specific subgenomes.

### SBP and ERF family TFs occupy chromatin interaction-linked gene regulation

Chromatin conformation regulates transcription by looping TF bound CREs to the promoters of distally located target genes, as observed in mammalian cell (*38, 59*) (**Fig. 5A**). To explore this mechanism in plants, we analyzed the enrichment of TFs at open chromatin interaction anchors. We found that binding sites of TF from the V-myb avian myeloblastosis viral oncogene homolog (MYB), DNA binding with one finger (Dof), ethylene responsive factor (ERF), GATA and SBP families were significantly enriched within the anchor region (**Fig. 5B**), with SBP and ERF binding motif-enriched loops exhibiting stronger interactions compared to others (**Fig. 5C)**.

**Fig. 5.**
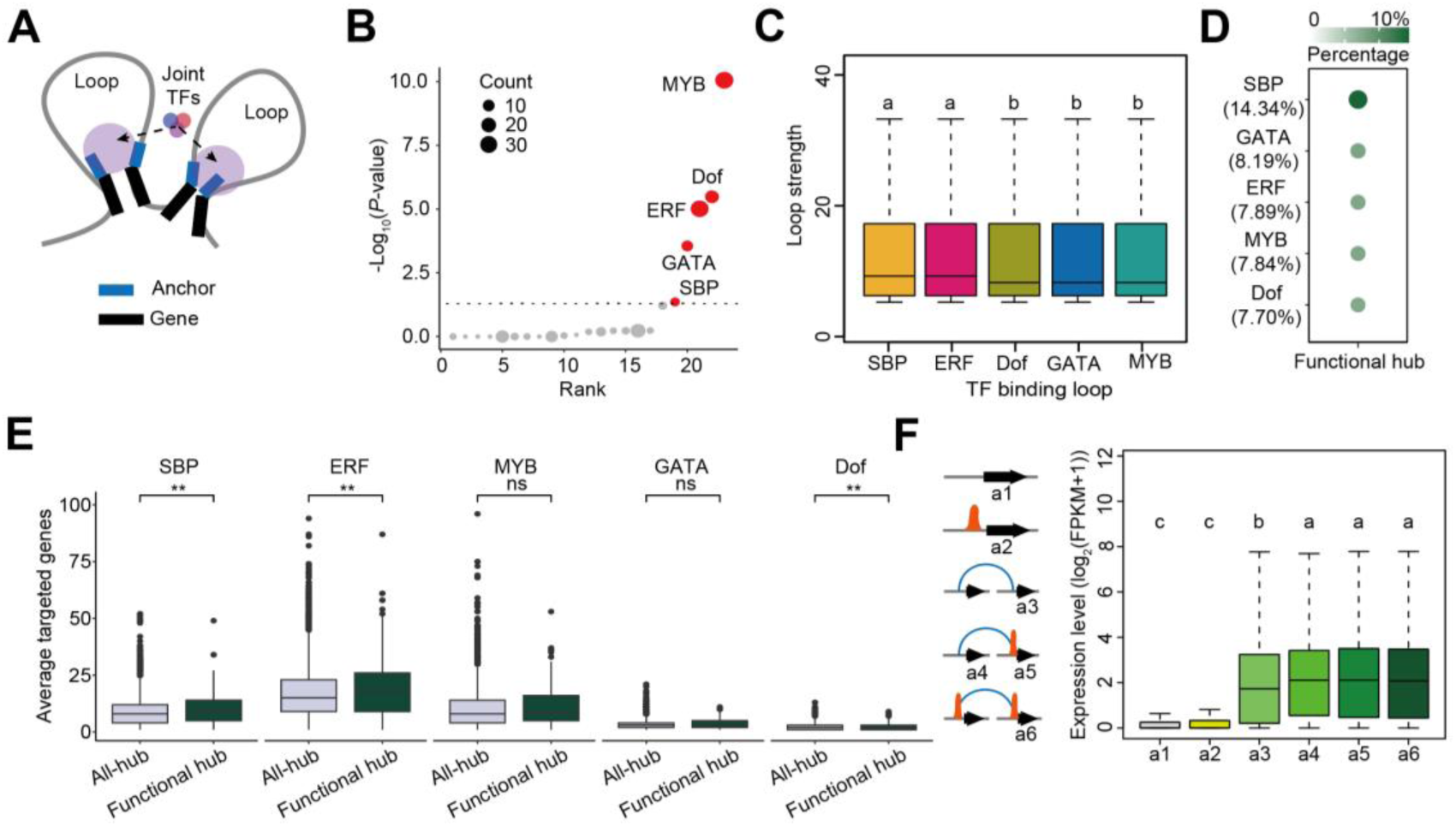
Identification of TFs enriched in the chromatin interaction regions. **A.** Schematic diagram illustrating joint TFs binding at the anchors of open chromatin interaction. **B.** TF family enrichment analysis in hub anchor regions (two-sided Fisher’s exact test). Significantly enriched TF families (*p*≤0.05) are highlighted. **C.** Boxplot showing loop strength across five TF families. The LSD multiple comparison test was used to assess significance. Different letters denote significant differences (*p*≤0.05). **D.** Percentage of TF binding sites enriched in functional hub anchor regions. Calculated as: TF binding site identified in functional hubs/TF binding sites identified in all hubs. **E.** Boxplot showing the average number of target genes (TF binding motifs in promoter regions) per hub anchor or functional hub anchor across five TF families. *, *p*≤0.05; **, *p*≤0.01 from Student’s *t*-test. **F.** Expression levels of genes categorized by chromatin interaction models and SBP occupancy. The LSD multiple comparison test was used to assess significance. Different letters indicate significant differences (*p*≤0.05).

To assess the biological significance of these TF binding sites-enriched anchors, we examined their presence in functional hubs (the hubs overlapped with previous defined functional regions). Notably, 14.34% of the SBP family TF binding sites were localized in functional regulatory regions, higher than other TF families (**Fig. 5D**). In addition, SBP and ERF-bound functional hub anchor linked to significantly more number of target genes compared to other hub anchors, indicating the importance of SBP and ERF targets in functional regions (**Fig. 5E**). Using published DAP-seq data for SBP, ERF, MYB, and Dof TF families, we found that the SBP and ERF DAP-seq peaks were more enriched in anchor regions (*60*) (**fig. S4A**), with SBP-bound loops showing enhanced interaction strength (**fig. S4B**). This suggest that SBP and ERF TFs are closely associated with chromatin interactions, emphasizing the importance of SBP-bound loops in transcriptional regulation.

Further analysis categorized genes based on SBP-associated chromatin interactions and occupancy (*44*) (**Fig. 5F**). Genes with SBP binding at one or both anchors (a4, a5, and a6) exhibited higher expression levels than those without SBP binding (a3) (**Fig. 5F**). However, basal genes not involved in chromatin interaction showed similar expression levels regardless of SBP binding (a1, a2), indicating that chromatin loops and SBP binding synergistically enhance transcription within chromatin loop.

In conclusion, we identified five TF family (MYB, Dof, ERF, GATA, and SBP) significantly enriched in open chromatin anchor regions, suggesting their role in the formation or functional regulation of chromatin interaction.

### TaSPL7/15-mediated chromatin interactions regulate photosynthesis and leaf development in wheat

To investigate the role of SBPs in stabilizing chromatin interactions, we analysed the CRISPR/Cas9-derived knockout mutant *Taspl7&15* (*61*) and conducted RNA-seq on *Taspl7&15* and wild-type (WT) counterpart, ZM7698. Transcriptomic analysis revealed strong correlations among replicates (**Fig. 6A**) and identified 1,917 down-regulated and 1,802 up-regulated genes in *Taspl7&15*. GO enrichment indicated pathways related to chloroplasts, leaf development, and carbohydrate transport were significantly affected, underscoring TaSPL7/15’s role in photosynthesis and leaf development (**Fig. 6B**).

**Fig. 6.**
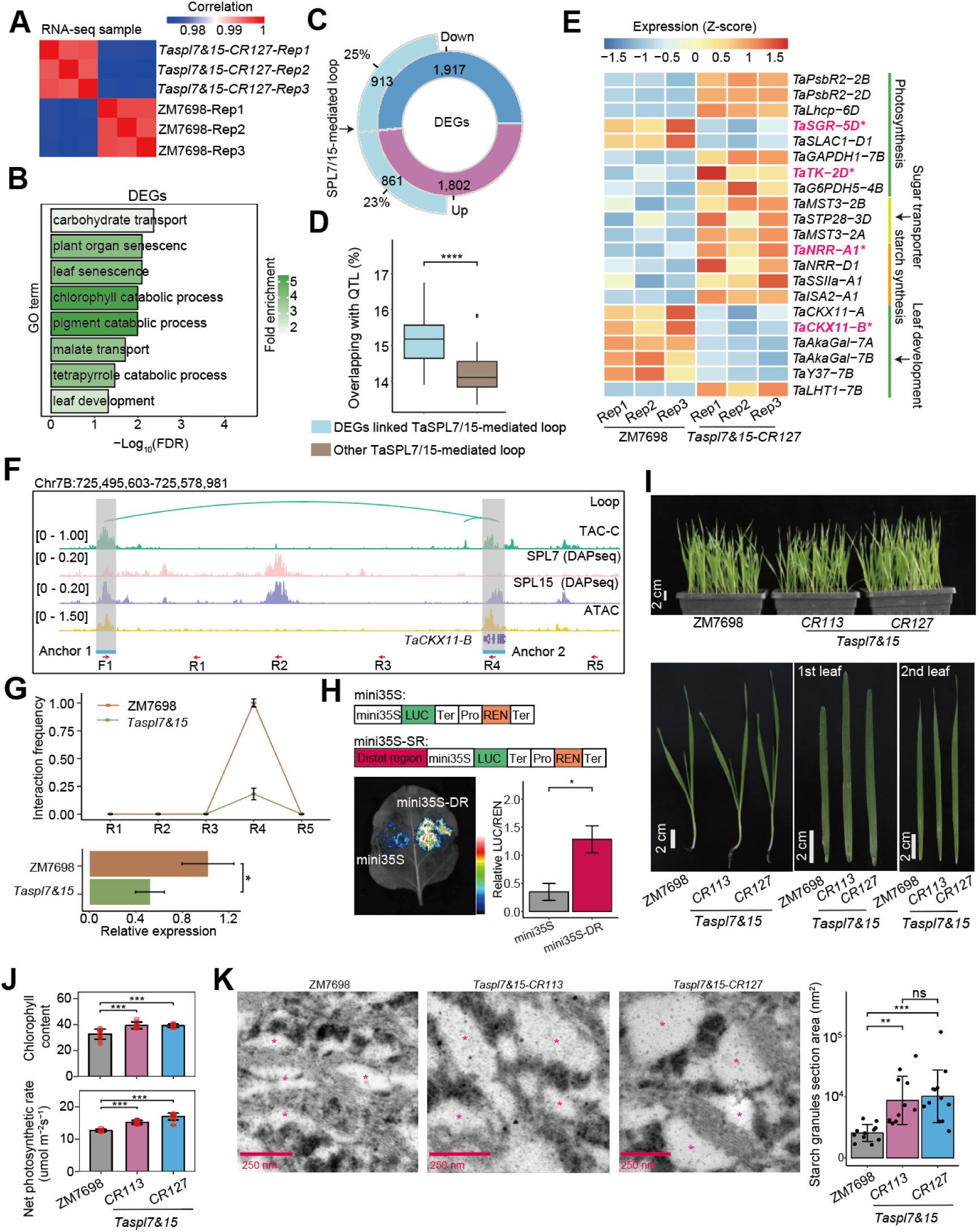
SPL-associated chromatin interactions in regulating photosynthetic energy metabolism. **A.** Clustering of samples for RNA-seq according to the person correlation coefficient of gene expression levels. **B.** GO enrichment analysis of DEGs in *Taspl7&15* vs ZM7698. **C.** Proportion of DEGs overlapped with TaSPL7/15-mediated chromatin loop linked genes. **D.** The percentage of overlapped interactions between TaSPL7/15-mediated loop linked DEGs and other TaSPL7/15-mediated loop. ***, *p*≤0.001 from Student’s *t*-test. **E.** Heatmap displaying the expression of key genes involved in photosynthesis and leaf development. Expression levels were normalized by *z*-score. The genes that are further validated in the following are highlighted. **F.** Browser screenshot showing chromatin loop interactions of *TaCKX11-B.* Profiles of TaSPL7/15 DAP-seq, ATAC, and chromatin interactions around *TaCKX11-B* and its distal regulatory site are shown, with anchor regions highlighted. The lower panel shows the locations of primer in TAC-C-qPCR experiments. **G.** The upper panel is the quantitative relative interacting frequency of loop regions determined by TAC-C-qPCR in ZM7698 and *Taspl7&15*. The lower panel is the quantitative relative expression of *TaCKX11-B* by RT-PCR assay in ZM7698 and *Taspl7&15*. Error bars: ±SD, *, *p*≤0.05 from Student’s *t*-test. **H.** Luciferase reporter assay validating the transcriptional regulatory role of the distal regions (DR) on *TaCKX11-B*. The distal anchor sequence was introduced into the proximal region of the *TaCKX11-B* promoter. *, *p*≤0.05 from Student’s *t*-test. **I.** Phenotypes of leaves of the wheat accession ZM7698 and two CRISPR-generated *Taspl7&15* mutant lines (*CR113* and *CR127*) at the seedling stage. **J.** Comparison of chlorophyll content, and net assimilation rates of carbon dioxide between ZM7698 and the two CRISPR lines of *Taspl7&15* mutant at the seedling stage. ***, *p*≤0.001 from Student’s *t*-test. **K.** Transmission electron microscopy (TEM) images of chloroplasts of for ZM7698 and *Taspl7&15* two mutant lines. Red asterisks indicate starch granules. The area of the starch granules section is calculated by ImageJ. **, *p*≤0.01; ***, *p*≤0.001 from Student’s *t*-test.

Approximately 48% of differentially expressed genes (DEGs) overlapped with TaSPL7/15-mediated chromatin loop-associated genes, suggesting that TaSPL7/15 maintains chromatin loops critical for gene expression (**Fig. 6C and table S5**). These loop-associated DEGs were significantly enriched in functional regulatory QTLs, highlighting their importance (**Fig. 6D**). Key loop-associated DEGs included chlorophyll a-b binding protein 5 (*TaLhcb5-6D*) (up-regulated), chlorophyll degradation gene *Stay-Green Rice* (*TaSGR-5D*) (down-regulated), and stomatal anion channel protein controlling stomatal closure gene *SLOW ANION CHANNEL-ASSOCIATED 1* (*TaSLAC1-D1*) (down-regulated), all influencing chlorophyll content and photosynthetic efficiency (**Fig. 6E, table S6**). Additionally, transketolase (TaTK-2D), positive regulator of starch accumulation in vegetative tissues CO_2_-responsive CCT protein (TaNRR-A1), were up-regulated, while negative regulators of leaf development, such as cytokinin oxidase/dehydrogenase 11 (TaCKX11-A) and (*TaCKX11-B*), were down-regulated.

To validate these chromatin interactions, we focused on *TaCKX11-B*, a gene previously linked to negative regulation of leaf size and photosynthetic efficiency in rice and wheat (*62*), (*56*). A 57-kb upstream interaction loop was identified in WT (ZM7698) but lost in *Taspl7&15* (**Fig. 6F**). TAC-C-qPCR confirmed the loss of this interaction loop in *Taspl7&15* (**Fig. 6G, upper**), and RT-qPCR demonstrated reduced *TaCKX11-B* expression (**Fig. 6G, lower**). Luciferase reporter assays validated the enhancer-like function of the distal region on *TaCKX11-B* (**Fig. 6H**). Similar loop disruption were observed for *TaSGR-5D*, *TaNRR-A1,* and *TaTK-2D* (**fig. S5A-C**).

Phenotypic analysis showed *Taspl7&15* mutants had larger leaf size, higher chlorophyll content, and improved photosynthetic efficiency (**Fig. 6I, J**). Liquid chromatography-mass spectrometry (LC-QQQ-MS) analysis revealed increased levels of photosynthetic intermediates, including pentose phosphate (D-Ribose 5-phosphate (R5P), ribulose-5-phosphate (Ru5P), D-Xylulose-5-phosphate (Xu5P), and D-Sedoheptulose-7-phosphate (S7P) in *Taspl7&15* compared to ZM7698 (**fig. S6**), indicating enhanced photosynthetic capacity. Notably, the absence of phosphoglycerate 3-phosphoglycerate (3PGA) and 2-phosphoglycerate (2PGA) suggested reduced photorespiratory flux, potentially boosting photosynthetic efficiency (**fig. S6**). Moreover, we observed larger starch granules in *Taspl7&15*’ leaves than in those of ZM9867 (**Fig. 6K**). Those phenotypic data fit well with the observed transcriptional change of above-mentioned genes in *Taspl7&15*.

In summary, TaSPL7/15 mediates chromatin interactions critical for regulating photosynthesis-related genes and leaf development. Loss of TaSPL7/15 disrupts these loops, altering gene expression and enhancing photosynthetic efficiency in wheat leaves.

## Discussion

Understanding the spatial organization of *cis*-regulatory elements (CREs) is essential for deciphering gene regulation (*63*), particularly in complex genomes like wheat, a hexaploid species (*29*). The three-dimensional structure of the genome plays a crucial role in how CREs, such as enhancers, interact with their target genes to control key biological processes (*44, 64*). Traditional methods like Hi-C, HiChIP, and OCEAN-C have advanced our understanding of chromatin interactions but have been less effective in capturing the fine-scale interactions between CREs and genes in wheat (*25, 28, 30*).

### TAC-C: A plant specialized fine-scale 3D chromatin organization study tool

Due to the higher spatial resolution of ATAC-seq compared to FAIRE-seq (*27*), we developed the TAC-C method by integrating ATAC-seq with *in situ* Hi-C, facilitating targeted capture of interactions within open chromatin regions. This advancement not only heightens spatial resolution but also reduces the sequencing depth required to yield meaningful data. This innovation holds particular significance for cost reduction, especially in the context of crops with large genomes like wheat. TAC-C excels in identifying active *cis*-regulatory elements and demonstrates better overlap with epigenetic modifications compared to other methods (**Fig. 1**). Using wheat as a model, TAC-C has proven its robustness in capturing intricate details of chromatin interactions that are crucial for understanding gene regulation in large-genome species. The method’s adaptation for plant samples, particularly those rich in sugars and polyphenols, further underscores its versatility and reliability. Unlike the recently developed ChIATAC technique (*18*), TAC-C omits the use of SDS in nuclei treatment (**Fig. 1**). SDS as a surfactant, has the potential to disrupt the binding between histones and DNA. This disruption facilitates Tn5 accessibility to closed chromatin regions. Consequently, compared to the ChIATAC, TAC-C is more similar to ATAC-seq.

Moreover, ChIATAC was only used in animal cells, with no reported application in plants, while TAC-C is a technology tailored specifically for plant systems. These makes TAC-C not only a powerful alternative but also a more specialized tool for studying the 3D genome organization in plants. The success of TAC-C in rice, wheat, maize, sorghum (**Fig. 2**) suggests its potential applicability across other complex genomes, providing a new avenue to explore the spatial organization and regulatory mechanisms underlying large-genome species.

### Conservation and biological importance of TAC-C identified chromatin interaction hub anchors

TAC-C identifies open chromatin regions frequently interacting with other regions, termed hub anchors (**Fig. 2**). Across different crops species, hub anchors exhibit common features such as low-frequency base variations and higher expression levels of their associated node genes (**Fig. 2**). Moreover, a significant overlap was observed between previously reported QTL, eQTL, and breeding selection intervals with chromatin interaction anchors, especially hub anchors (**Fig. 3**). This underscores the functional importance of open chromatin interactions in regulating agronomic traits. GWAS has identified numerous QTLs in crops with large genomes like maize and wheat (*65, 66*), often located in intergenic regions, complicating their functional interpretation. Chromatin interactions can be used as physical evidence to connect GWAS-identified SNPs to their target genes, offering insights into trait-associated elements. For example, our study revealed chromatin interactions between *TaNAL1-5A-like* and its eQTL loci (**Fig. 3**). This research advances our understanding and paves the way for precision breeding using advanced editing technique.

Understanding the differences in transcriptional regulation between C3 and C4 plants, and uncovering the related *cis*- and *trans*-regulatory evolution, has long been a key goal in the field of plant biology (*67*). Previous studies have identified a limited set of specific ACR and TF-mediated regulatory mechanisms that govern the expression of C4 genes (*68–71*). Our study reveals differences in open chromatin interactions between C4 and C3 plants, highlighting a more frequent interaction of C4-speicifc genes (**Fig. 2**), such as *PEPC*, with other regions in C4 plants. This discovery offers new insights into the regulatory changes that occurred during the evolution of C4 photosynthesis, providing a fresh perspective on the evolution of C4 plants.

### Chromatin spatial organization contributes to subgenome gene expression bias

Previous research has demonstrated that differences in epigenetic modifications, transposon insertions, and TF binding motifs significantly influence homoeolog expression bias in wheat (*41, 58, 72, 73*). Our TAC-C data support these findings, revealing extensive subgenome-specific PPIs (**Fig. 4**). Notably, the D subgenome exhibited more chromatin interactions compared to the A and B subgenomes (**Fig. 4**), possibly due to the latter undergoing epigenetic homogenization over time (*74*). In contrast, the D subgenome, with its shorter evolutionary history, may retain more diploid-like epigenetic features.

Our results also show an enrichment of transposons in subgenome-specific interaction anchors (**Fig. 4**), suggesting that some transposons have evolved into regulatory elements, influencing subgenome-specific gene expression through chromatin looping (*60, 73, 75*). While previous studies focused on genetic sequence variations and epigenetic modifications (*41, 60, 76*), our research highlights the importance of distal chromatin interactions in shaping subgenome bias (**Fig. 4**). We propose that sequence variations and transposon insertions near homoeologous genes modify the chromatin environment, leading to biased gene expression and reinforcing subgenome imbalances. This asymmetric chromatin interaction offers new insights into the regulation of subgenome bias in hexaploid wheat and presents opportunities for developing precise gene regulation strategies at the subgenome level.

### SBPs in chromatin organization and wheat development

While CTCF/cohesin complexes control chromatin loops in animals (*35, 36*), and TCPs shape chromatin structure in *Marchantia* (*40*), the processes governing chromatin loops in higher plants like wheat are not well understood (*77*). TaSPL7/15 participate in wheat vernalization process by directly anchoring the *VERNALIZATION1* (*VRN1*) promoter and 30Kb upstream regulatory elements of *VRN3* (*78*), yet their broader role in wheat 3D chromatin structure has been unexplored. Our findings reveal an enrichment of SBP and ERF TF binding motifs in chromatin’s open anchor regions (**Fig. 5**), indicating the potential role in maintaining wheat chromatin organization. In *Taspl7&15* mutants, a substantial overlap was observed between differentially expressed genes and TaSPL7/15-mediated loop linked genes (**Fig. 6**), particularly those involved in photosynthesis, starch synthesis, sugar transport, and leaf development (**Fig. 6**), leading to increased photosynthetic efficiency in *Taspl7&15*. Specifically, the loss of distal regulatory interaction for *TaCKX11-A*, *TaCKX11-A*, *TaSGR*-*5D*, *TaTK-2D* and *TaISA2* resulted in longer leaves, reduced chlorophyll degradation, enhancing photosynthesis efficiency and increased photosynthetic products (**Fig. 6**). Additionally, SPLs are quantitatively regulated by miRNA156s, conserved microRNAs associated with plant developmental ages across species (*79, 80*), suggesting that 3D chromatin organization may undergo SPL-mediated change as plants mature, contributing to developmental morphology.

## Methods

### Plant materials and growth conditions

The *O. sativa*, *S. bicolor*, *Z. mays* and *T. aestivum* (Chinese Spring) were used in this study. For the *Taspl7&15* CRISPR lines which we got from previous study (*61*). The seeds were surface sterilized in 2% NaClO for 20 minutes, then rinsed overnight with flowing water. The seeds were germinated and then transferred to soil, and grown in the greenhouse at 22°C/20°C day/night, under long day conditions (16h light/8h dark). When the material grows to one leaf and one heart period, the leaves were sampled for the RNA-seq (three replicates), and TAC-C experiments (two replicates).

### TAC-C library construction

TAC-C experiments require three processions, including:

Plant tissue was cut into approximately 0.5 square centimeter pieces, and then fixed using Nuclei Isolation Buffer (NIB) containing 4% formaldehyde (20mM Hepes, 250mM Sucrose, 1mM MgCl_2_, 5mM KCl, 1mM DTT, 1% Protease Inhibitor Cocktail for Plants, 0.25% Triton X-100, 40% glycerol, 4% formaldehyde). The reaction was quenched by adding 0.125 M glycine and incubated on ice for 15 minutes. The fixed plant tissue was washed three times with NIB (20mM Hepes, 250mM Sucrose, 1mM MgCl_2_, 5mM KCl, 1mM DTT, 1% Protease Inhibitor Cocktail for Plants, 0.25% Triton X-100, 40% glycerol). After removing excess liquid from the surface using absorbent paper, nuclear extraction could be performed directly, or the samples could be stored at −80℃.

The processed plant tissue, subjected to liquid nitrogen grinding, was transferred to a 15mL tube, and the pellets were then resuspended using NIB. The suspension was incubated with shaking on ice for 10 minutes. Subsequently, the suspension was successively filtered through 10μm and 40μm strainers, followed by centrifugation at 1260g at 4℃ for 5 minutes. After discarding the supernatant, the pellets were washed twice with NIB and then washed twice with 1.2×NEBuffer r2.1 (NEB; B6002S). Nuclei were obtained after centrifugation at 1260g at 4℃ for 5 minutes with the removal of the supernatant. The isolated nuclei can be used for TAC-C library preparation or stored at −80℃.

Using 10,000 to 50,000 nuclei as the starting material for TAC-C library construction. The nuclei were resuspended in 500μL 1.2× NEBuffer r2.1, and 40μL DpnII (NEB; R0543L) was added. The reaction was carried out overnight at 37℃ with constant agitation at 900rpm in a metal bath. Subsequently, the reaction was terminated by incubating at 65℃ for 20 minutes. After that, 37.5μL Biotin-14-dCTP (Invitrogen; 19518018), 1.5μL 10mM dATP (NEB; N0446S), 1.5μL 10mM dTTP (NEB; N0446S), 1.5μL 10mM dGTP (NEB; N0446S), and 10μL DNA Polymerase I, Large (Klenow) Fragment (NEB; M0210L) were added, and the reaction was carried out at 37℃ for 1h in a metal bath to complete biotin labeling. Following this, 598μL 2× Rapid Ligation Buffer and 6 μL T4 DNA ligase (Enzymatics; L6030-HC-L) were added, and the reaction was carried out at 20℃ for 4h to complete the *in situ* ligation. The reaction mixture was centrifuged at 5000g for 5 minutes, the supernatant was discarded, and 1μL Exonuclease III (NEB; M0206), 10μL 10×NEBuffer1 (NEB; M0206), and 89μL Nuclease-Free Water were added. The reaction was carried out at 37℃ for 20 minutes, followed by 72℃ for 30 minutes to terminate the reaction. After washing once with 500μL 1×TD buffer (10mM Tris-HCl, pH 7.4, 5mM MgCl_2_, and 0.25% Dimethyl Formamide), the samples were centrifuged at 1260g for 5 minutes, and the supernatant was discarded. Then, 50μL 2×TD Buffer (20mM Tris-HCl, pH 7.4, 10mM MgCl_2_, 0.5% Dimethyl Formamide) and 8μL TTE Mix V50 (Vazyme; TD711-02), 42μL Nuclease-Free Water were added, and the reaction was carried out at 900rpm, 37℃ for 1h in a metal bath. After the reaction, the DNA was purified using the MinElute PCR Purification Kit (Qiagen; 28004), washed with 100μL Buffer EB (Qiagen; 19086), and then incubated with 40μL Dynabeads MyOne Streptavidin C1 (Invitrogen; 65001) at room temperature for 20 minutes. The beads were washed twice with Binding and Washing Buffer (5mM Tris-HCl, pH 7.5, 0.5mM EDTA, 1M NaCl). Finally, the DNA was amplified by adding 50μL TruePrep CUT&Tag Amplification Mix (Vazyme; TD612-01), 40μL Nuclease-Free Water, 5μL 20uM N7 index primer (TGTGAGCCAAGGAGTTGTTGTCTTCNNNNNNNNNNGTCTCGTGGGCTCGG), and 5μL 20uM N5 primer (/5Phos/GAACGACATGGCTACGATCCGACTTTCGTCGGCAGCGTC). The amplification conditions were as follows: 72℃ for 5 minutes, 98℃ for 30 seconds; 98℃ for 10 seconds, 63℃ for 30 seconds, 72℃ for 1 minute, 10 cycles; 72℃ for 5 minutes; hold at 4℃. The PCR products were purified using 60μL Ampure beads (Beckman Coulter; A63881), incubated for 5 minutes, placed on a magnetic stand for 5 minutes, and then the supernatant was transferred to a new 1.5mL tube. After incubating with 20μL Ampure beads for 5 minutes and placing on a magnetic stand for 5 minutes, the supernatant was discarded. The beads were washed twice with 80% ethanol, and after the beads were air-dried, the DNA was recovered using Buffer EB. This is the TAC-C library. The TAC-C library was sequenced using MGI 2000 (2×100bp). The relevant chemicals and enzymes are listed in **table S7**.

### RNA-seq library construction

Total RNAs were extracted using Ultrapure RNA kit (CWBIO; CW0581M) according to manufacturer instructions. The products were sequenced using an Illumina NovaSeq6000 by Gene Denovo Biotechnology and RNA-seq data were obtained from three biological replicates.

### TAC-C data processing

Initially, the raw reads underwent trimming using the hicup_truncater v0.5.9 tool (*81*) to identify and eliminate the ligated junction points within the reads, specifically removing the sequences after the recognition site of restriction enzyme. Subsequently, the trimmed reads were mapped to the reference genome: *S. bicolor* (*82*), *Z. mays* (*83*), *O. sativa* (*84*) and *T. aestivum* (*85*) through the bwa v0.7.17 software (*86*), with reads having a mapping quality (MAPQ) score<1 being discarded to ensure mapping accuracy. The Hi-C-Pro v2.10.0 (*87*) tool was employed to filter out Dangling End Pairs, Re-ligation Pairs, Self-cycle Pairs, and Dumped Pairs. Following the removal of PCR duplicates, the remaining valid read pairs were used for calling peaks and loops. The calling of peaks from the remaining valid read pairs was performed using MACS2 (*88*) the parameter “-q 0.01 -f BAM --nomodel --extsize 150 --shift −75 --call-summits --nolambda”, The overlapping peaks between two biological replicates were used as an anchor to identify open chromatin interactions by hichipper (http://aryeelab.org/hichipper) with default setting in two biological replicates. Finally, the intrachromosomal loop from two biological replicates were merged by hicMergeLoops from HiCExplorer software (*89*) with the parameter “-r 10000”. TAC-C interaction heatmaps were produced with HiCPlotter v0.8.1 at 1-Mb resolution (*90*).

### OCEAN-C and Hi-C data processing

We downloaded the OCEAN-C and Hi-C data of Chinese Spring from the NGDC BioProject database (accession number CRA003731) and NCBI (accession number GSE133885) (*25, 28*). Reads were aligned to the IWGSC RefSeq v2.1 reference genome. The data analysis procedures are the same with the analysis for TAC-C data. In particular, for the Hi-C intrachromosomal loops, we directly use the published results of previous study (*30*).

### RNA-seq data analysis

Low-quality reads and adapter sequences were removed using Cutadapt (*91*) and Trimmomatic (*92*), respectively. Retained clean RNA-seq reads were mapped to the same version reference genome with TAC-C by using STAR(*93*) (version 2.5.4b) with default parameters: SortedByCoordinate --quantMode TranscriptomeSAM GeneCounts, and accurate transcript was quantified using RSEM (*94*) (version 1.3.0). In addition, RSEM was also used to analyze the mRNA expression level by calculating Fragments per Kilobase of exon model per Million (FPKM) mapped reads. The RNA-seq reads counts of DEGs were filtered with |log_2_(fold change)|>0 and FDR<0.05 by using R packages edgeR (*95*).

### Gene ontology analysis

The GO functional enrichment was performed using the find_enrichment.py from goatools software (*96*).

### ChIP-seq and ATAC-seq data analysis

Sequencing reads of H3K4me3, H3K9ac, H3K27ac, H3K27me3, H3K9me3, H3K36me3 and ATAC-seq of Chinese Spring were obtained from previous studies (*28, 41*). ChIP-seq data are available in NCBI GEO under accession number GSE121903 and ATAC-seq data are available in NGDC GSA under accession number CRA003731. Sequencing reads were cleaned by removing bases that had low quality scores (<20) and removing sequencing adaptors by Cutadapt (*91*) and filtering out short reads by Trimmomatic (*92*). The reads of ChIP-seq and ATAC-seq were mapped to the IWGSC RefSeq v2.1 reference genome using Bowtie2 (*97*) with default setting. The concordantly mapped reads (MAPQ>10) were kept and PCR duplication was further removed with Picard (https://broadinstitute.github.io/picard/). Peaks of ChIP-seq libraries were detected using MACS2 (2.1.2) with the parameter “-f BAM –nomodel - q 0.05”. The peaks of ATAC-seq were identified using MACS2 with the parameter “-q 0.05 -f BAM --nomodel --extsize 200 --shift −100”.

### DAP-seq data analysis

For SPL7A, SPL7B, SPL7D, SPL15A, SPL15B and SPL15D DAP-seq data analysis, the raw sequencing reads (from the NCBI accession number PRJNA779959) (*61*) were cleaned by removing bases that had low quality scores (<20) and removing sequencing adaptors by Cutadapt (*91*), and filtering out short reads by Trimmomatic (*92*). As a result, about 5 million reads with a MAPQ score>20 was obtained for further analyses. The cleaned reads were mapped to the IWGSC RefSeq v2.1 reference genome using Bowtie2 (*97*) (version 2.4.4) with default settings. MACS2 (*88*) was used for peak calling of the ChIP-Seq datasets, with options: -g 7.83e8 --call-summits --bdg -q 0.01 - -nomodel. Typical DAP-seq loci were plotted using Integrative Genomics Viewer solftware (*98*).

For ERF, Dof, MYB and other SPL DAP-seq peak were download from GSE192815 and GSE188699 (*60, 61, 73*). The overlap of DAP-seq peak with anchor were calculated using BEDTools (*99*) with parameter: intersect -wa -wb.

### Identification of homoeolog PPIs

The interspecies syntenic genes and homologous genes were download from TGT (http://wheat.cau.edu.cn/TGT/) across four species (*100*). PPIs whose anchors on one side were homologous genes between two species and anchors on another side was also homologous genes were defined as homologous PPIs, and the corresponding genes are defined as homologous PPI genes. The PPI homologous in *S. bicolor* and *Z. mays* (*83*) were defined as C4 conserved PPIs; PPI homologous in *O. sativa* and *T. aestivum* were defined as C3 conserved PPIs.

High-confidence gene models from the IWGSC (version 1.0) were used for defining triad genes. Homoeolog genes between each pair of A, B, and D subgenomes were identified as previously described (*58*). The Gene ID from IWGSC (version 1.0) were converted to IWGSC (version 2.1) by TGT (http://wheat.cau.edu.cn/TGT/) (*100*). Then a total 18,333 homoeolog groups with only one gene copy in each subgenome were defined as triads. PPIs whose anchors on one side were homoeolog genes between A, B, or D subgenomes and anchors on another side was also homoeolog genes were defined as homoeolog PPIs, and the corresponding genes are defined as homoeolog PPI genes.

### TF Motif analysis

We use Homer (*101*) to find enrichment of sequence motifs. Sequences in TAC-C anchor in were used as input. The “Known Results” were used as final results. The enriched TF were further used for TF family enrichment analysis. Enrichment analysis was done using the R package clusterProlifer (v3.16.1) (*102*), with a threshold *P*_adj_ < 0.05.

### RT- qPCR and TAC-C-qPCR

Total RNA was extracted using HiPure Plant RNA Mini Kit according to the manufacturer’s instructions (Magen; R4111-02). First-strand cDNA was synthesized from 2μg of DNase I-treated total RNA using the TransScript First Strand cDNA Synthesis SuperMix Kit (TransGen; AT301-02). RT-qPCR was performed using the ChamQ Universal SYBR qPCR Master Mix (Vazyme; Q711-02) by QuantStudio5 (Applied biosystems). The expression of interested genes was normalized to Tubulin for calibration, and the relative expression level is calculated via the 2^-ΔΔCT^ analysis method (*103*). Primers used for RT-qPCR are listed in table S8.

TAC-C-qPCR was performed to calculate the relative interaction frequencies between the two regions, as described previously TAC-C-qPCR with minor modifications (*104*). A genomic region without DpnII digestion was amplified as a loading control to normalize the DNA concentrations of different samples. Primers used for TAC-C-qPCR are listed in table S8.

### Luciferase reporter assay

To generate the p*TaNAL1-like-5A*:LUC construct, we amplified 3-Kb promoter fragments upstream of *TaNAL1-like-5A* from CS and ligated them with the transient expression vector CP461-LUC, which was constructed as the reporter vector. The p35S-*Traes5A02G000700*-GFP construct was used as effector. Then, p*TaNAL1-like-5A*:LUC and p35S:*Traes5A02G000700*-GFP were transformed into *A. tumefaciens* strain GV3101. The p*TaNAL1-like-5A*:LUC and p35S-pro:GFP were co-transformed as controls.

To generate *PR*-mini35Spro:LUC construct, the genomic sequence of distal regulatory region was amplified and fused in-frame with the pMY155-mini35S vector to generate the reporter construct. Then, mini35Spro:LUC (as control) and *PR*-mini35Spro:LUC were transformed into *A. tumefaciens* strain GV3101.

Then these strains were injected into *N. benthamiana* leaves (6∼8 leaf stage) in different combinations with p19, which was used to suppress RNA silencing. Dual luciferase assay reagents (Promega; VPE1910) with the Renilla luciferase gene as an internal control were used for luciferase imaging. The Dual-Luciferase Reporter Assay System kit (Promega; E2940) was used to quantify fluorescence signals. Relative LUC activity was calculated by the ratio of LUC/REN. The relevant primers are listed in **table S8**.

### Measurement of photosynthetic capacity, chlorophyll content, and leaf area

The measurement of photosynthetic capacity was conducted on the *Taspl7&15*-*CR* lines. The penultimate new leaves fully expanded were selected for the measurement of net photosynthesis rate with a portable gas-exchange and fluorescence system (GFS- 3000, Walz, Germany). The environmental parameters were set as follows: an air-flow of 400 mmol s^−1^ through the chamber, a leaf temperature of 25°C, a CO_2_ concentration of 400 μmol mol^−1^.

Chlorophyll content was measured with the same penultimate new fully expanded leaves at room temperature with a TYS-B Chlorophyll meter (Zhejiang TP, China).

Leaf area was measured with the same penultimate new fully expanded leaves at room temperature with a Yaxin-1241 portable Leaf Area Meter (Beijing Yaxinliyi Science and Technology Co. Ltd., China).

### Liquid Chromatography-Mass Spectrometry analysis

The endogenous levels of R5P/Ru5P/Xu5P, 3PGA/2PGA, E4P, DHPA, S7P, F6P/G1P and G6P were determined with an ultra-performance liquid chromatography-tandem mass spectrometry analytical platform consisting of an Agilent 1290 Infinity liquid chromatography pump and a 6495 triple quadrupole mass spectrometer (Agilent Technology Co. Ltd., America), following the protocol described in previous study (*105*).

### Ultrathin sections and scanning electron microscopy

Ultrathin (70nm) sections of ZM9867 and *Taspl7&15*-*CR* leaves were prepared using a Leica EM UC7 ultramicrotome (Leica, Vienna, Austria), as previously described (*106*). Sections were placed on formvar-coated copper grids and poststained with uranyl acetate and lead citrate and observed under a HT-7700 transmission electron microscope (Hitachi, Tokyo, Japan) operated at 80 kV.

### Statistics and data visualization

If not specified, R (https://cran.r-project.org/; version 4.3.2) was used to compute statistics and generate plots. For the two groups’ comparison of data, the Student’s *t*-test was used such as Fig. 2, 3, 4, 5, 6, 7 and fig. S2, 3 and 4. For three or more independent groups comparison of data, Fisher’s Least Significant Difference was used, such as Fig. 3B, 5C and 5F. Pearson correlation was used in Fig. S1A. For enrichment analysis, Fisher’s exact test was used, such as Fig. 5B, 6B, fig. S2H, fig. S3C and S3D. The Integrative Genomics Viewer (IGV) was used for the visual exploration of genomic data, such as Fig. 1I, 2I, 3F, 4I, 6F and fig. S5A-C

## Data and materials availability

Sequencing data of TAC-C, and RNA-seq are available at Genome Sequence Archive of the National Genomics Data Center, China National Center (<u>PRJCA029954</u>) and are publicly accessible at https://ngdc.cncb.ac.cn/gsa. Code used for all processing and analysis is available in GitHub (https://github.com/ZhangZhaoheng24/TAC-C).

## Acknowledgements

We thank Prof. Yiping. Tong (Institute of Genetics and Developmental Biology, Chinese Academy of Sciences) for the *Taspl7&15*-CR mutants and Dr. Chunyan Zhang (Plant Science Facility of the Institute of Botany, Chinese Academy of Sciences) for the help on the measurement of photosynthetic capacity, Prof. Yue Zhou (Peking University) for the comments on the manuscript.

## Author contributions

J.X. designed and supervised the research. J.X., Z.-H.Z., X.-L.L. and J.-M.K. wrote the manuscript. J.-M.K. developed and tested the TAC-C technology. X.-L.L. did the sample collection, mutants phenotype and physiological experiments. J.-M.K. and Y.-S. did TAC-C experiments. X.-M.L. did transcription regulation assay. Z.-H.Z., F.-Y.L., P.Z. and Y.-J.L. performed data analysis. X.-Y.L., Y.-Y.L. and W.-D.W. helped for the photosynthetic phenotypic analysis. C.-M.L., S.-B.X. and X.L. revised the manuscript. Z.-H.Z., X.-L.L. and J.X. prepared all the figures. All authors discussed the results and commented on the manuscript.

## Supplemental figures and legends

**Fig. S1.**
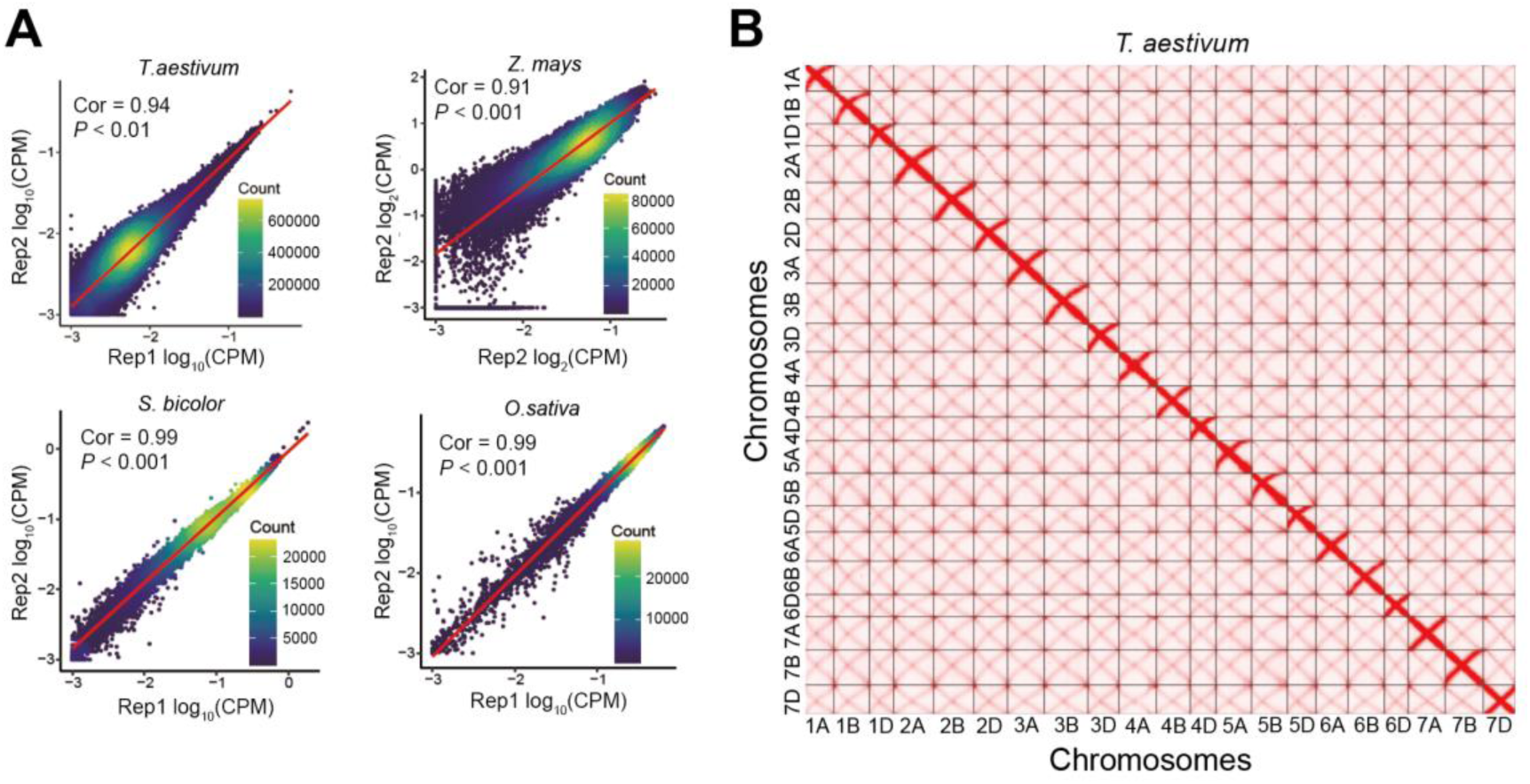
Reproducibility and contact matrix of TAC-C. **A.** Scatter plots showing the reproducibility between two TAC-C replicates of *O. sativa*, *S. bicolor, Z. mays* and *T. aestivum*. **B,** TAC-C contact matrix of the hexaploid wheat across 21 chromosomes

**Fig. S2.**
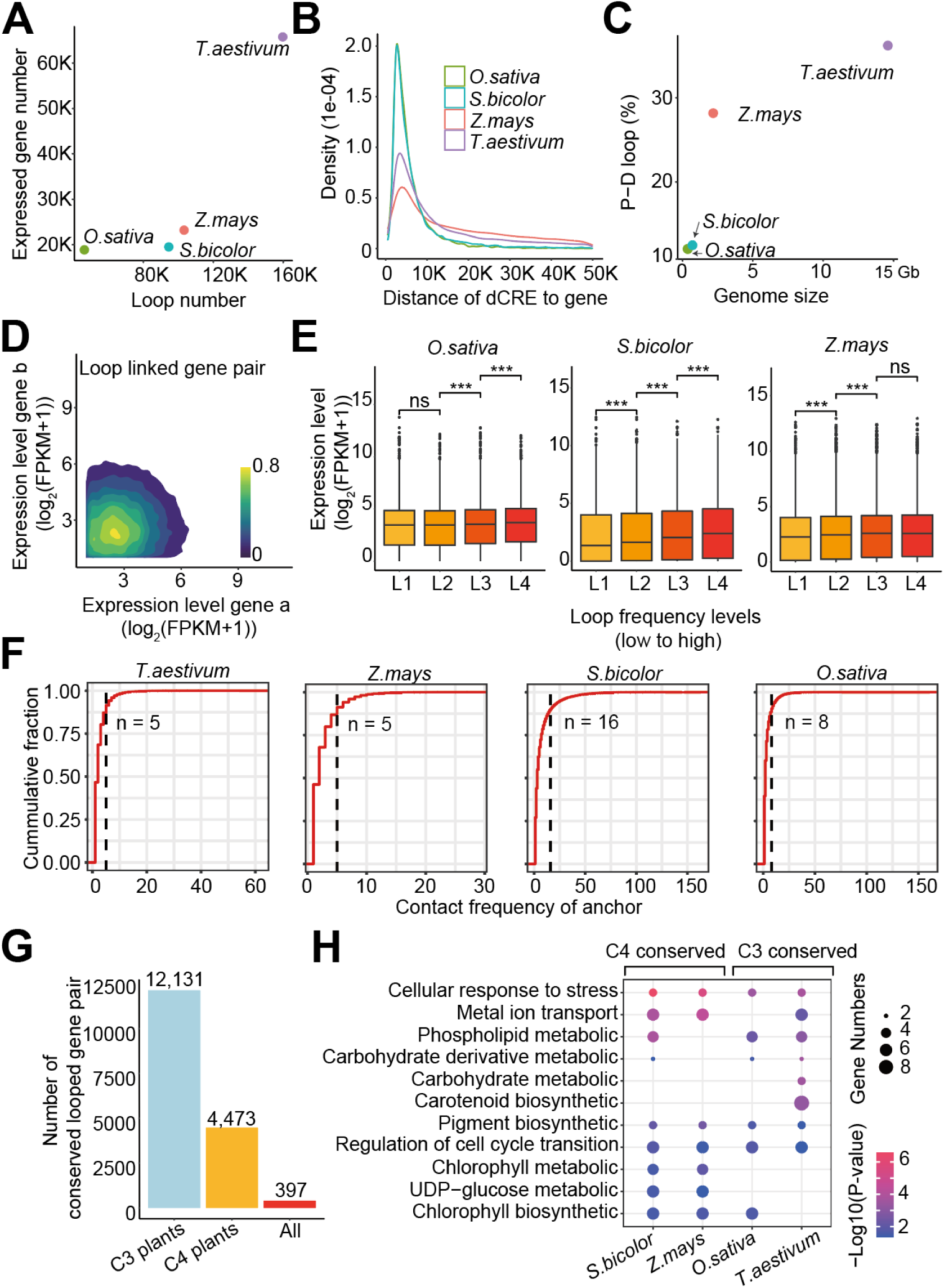
Characteristics of chromatin interactions in crop species. **A.** Correlation between loop counts and expressed gene across the four crop species. Genes with TPM>0.5 were considered expressed. **B.** Distribution of the distance between dCRE and transcription start site (TSS) of genes across the four crop species. **C.** Association between the proportion of PDI loop and genome size across the four crop species. **D.** Contour plot showing log-transformed FPKM values for gene pairs connected by TAC-C loops. **E.** Expression levels (log_2_(FPKM+1)) of genes with varying loop frequencies in *O. sativa*, *S. bicolor* and *Z. mays*. Genes were classified into four groups based on loop frequency. ***, *p*<0.001, ns, *p*>0.05 from Student’s *t*-test. **F.** Cumulative fraction of contact frequency for anchors in the four crop species. Dashed lines indicate the threshold, with anchors exceeding the frequency defined as hub anchors. **G.** Number of conserved loop-linked gene pairs in C4 plants (*S. bicolor* and *Z. mays*) and C3 plants (*O. sativa* and *T. aestivum*). **H.** GO terms for conserved genes linked by loops in C4 or C3 plants.

**Fig. S3.**
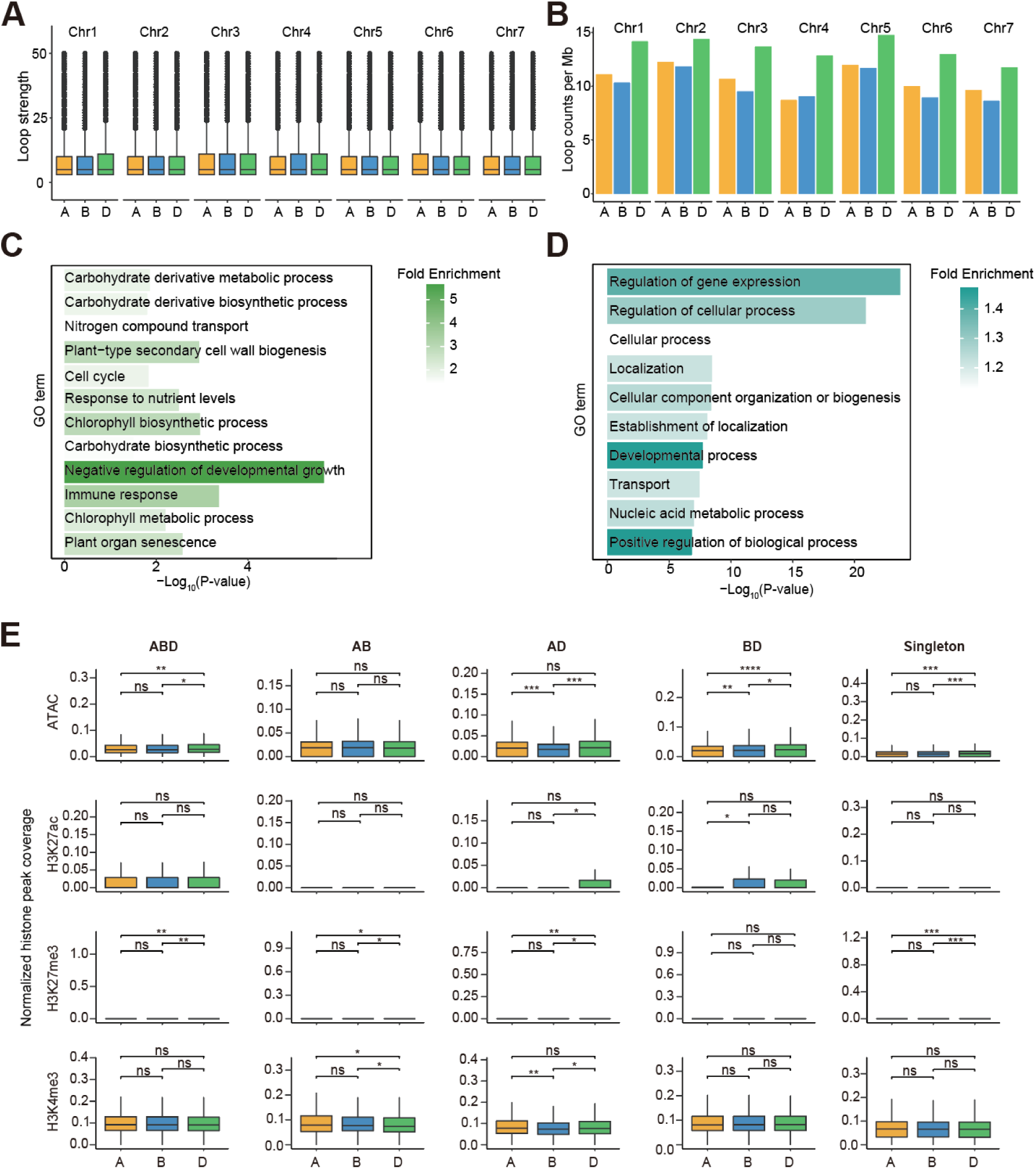
Open chromatin loops across the A, B and D subgenomes. **A.** Boxplot showing loop strength across the 21 wheat chromosomes. **B.** Loop counts per Mb across the 21 wheat chromosomes. **C.** GO enrichment analysis of genes linked to ABD homoeolog PPIs. **D.** GO enrichment analysis of genes linked to singleton PPIs. **E.** Coverage of ATAC, H3K27ac, H3K27me3, and H3K4me3 peaks in the 5 kb flanking regions of PPI-linked genes at different homoeolog levels in subgenomes A, B, and D. *, *p*≤0.05; **, *p*≤0.01; ***, *p*≤0.001 from Student’s *t*-test.

**Fig. S4.**
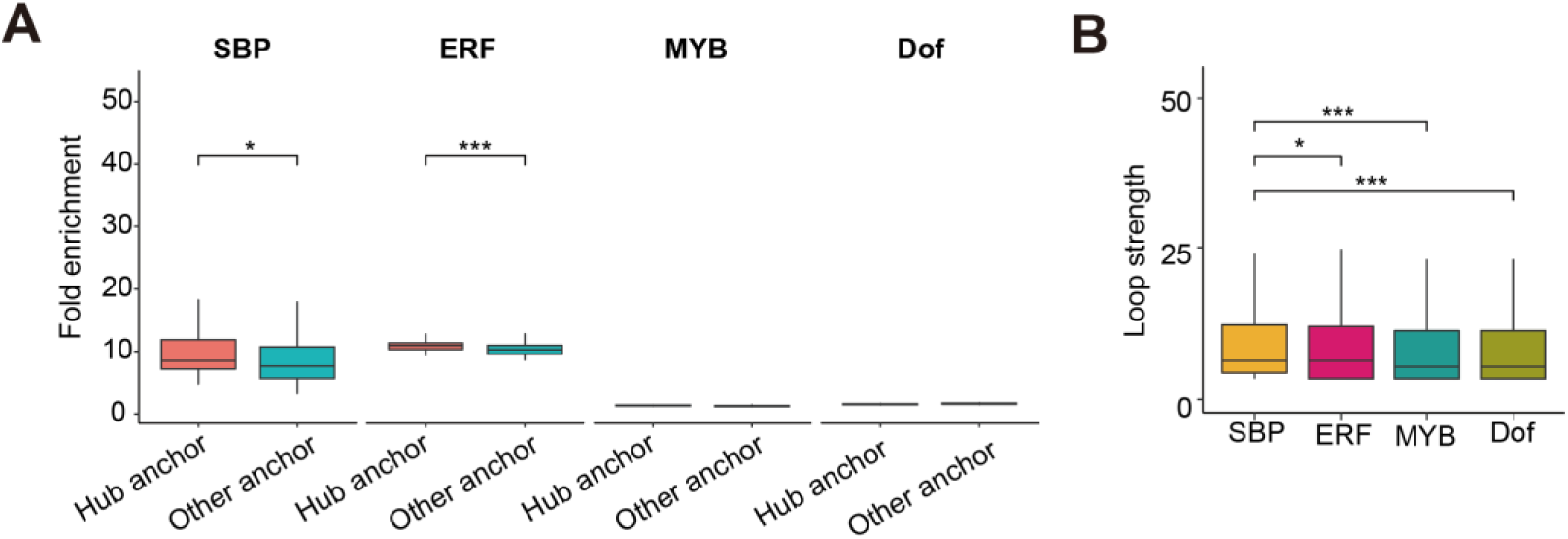
Difference in TF-associated chromatin topology. **A.** The fold enrichment (compare to random regions) of overlapped interactions between hub anchor, other anchor and random regions with SBP, ERF, MYB and Dof DAP-seq peaks. To minimize false positive errors, the 100 permutation tests were conducted in comparison. In each permutation test, 2000 accessions were randomly sampled from each of the different compared groups. *, *p*≤0.05; ***, *p*≤0.001 from Student’s *t*-test. **B.** The boxplot shown the strength of loop containing five TF families DAP-seq peak. *, *p*≤0.05; ***, *p*≤0.001 from Student’s *t*-test.

**Fig. S5.**
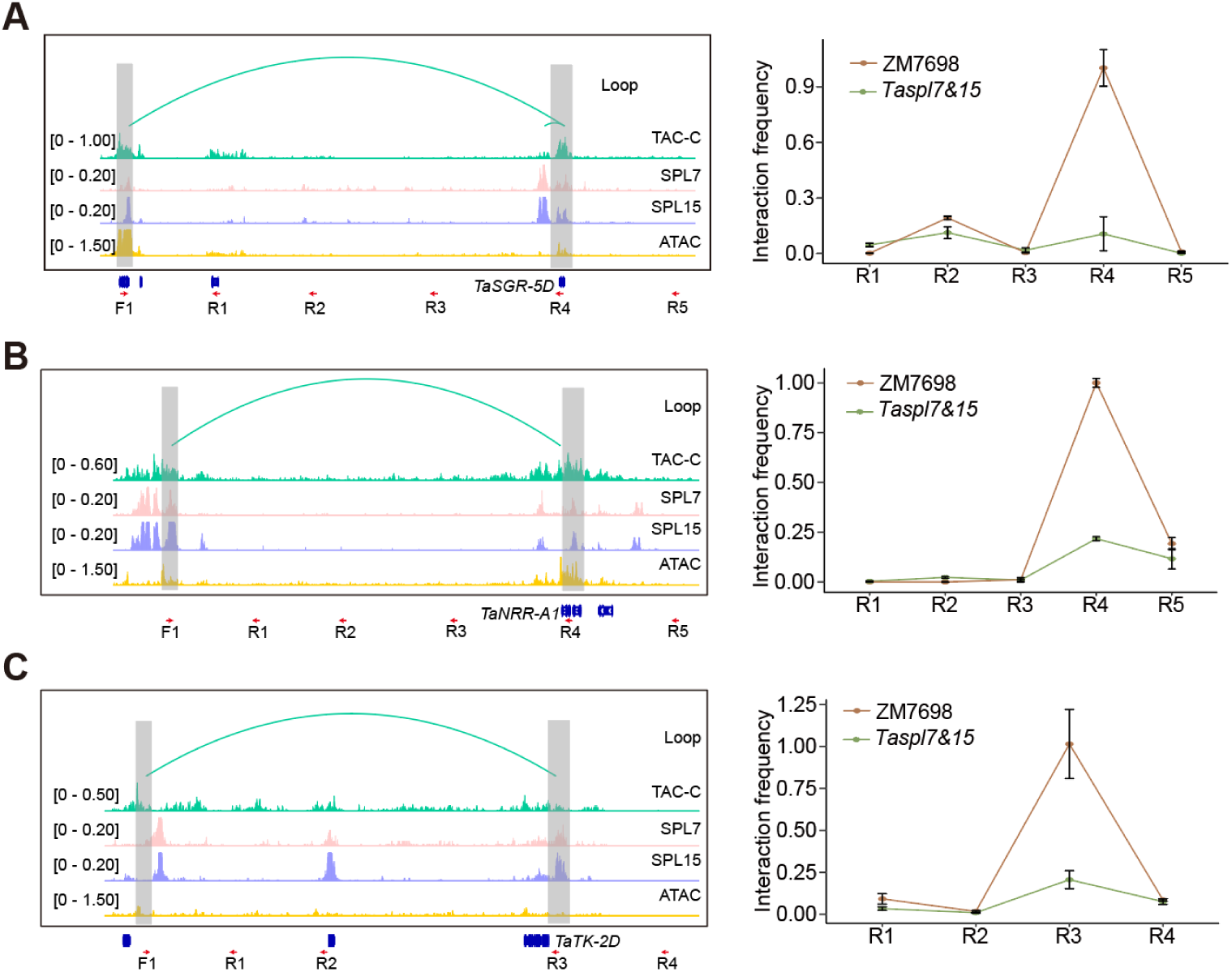
Examples of loops lost in *Taspl7&15* compared to ZM7698. **A-C.** The left panel is the quantitative relative interacting frequency of loop regions determined by TAC-C-qPCR in ZM7698 and *Taspl7&15*. The right panel is the quantitative relative expression of *TaCKX11-B* by RT-PCR assay in ZM7698 and *Taspl7&15*. Error bars: ±SD.

**Fig. S6.**
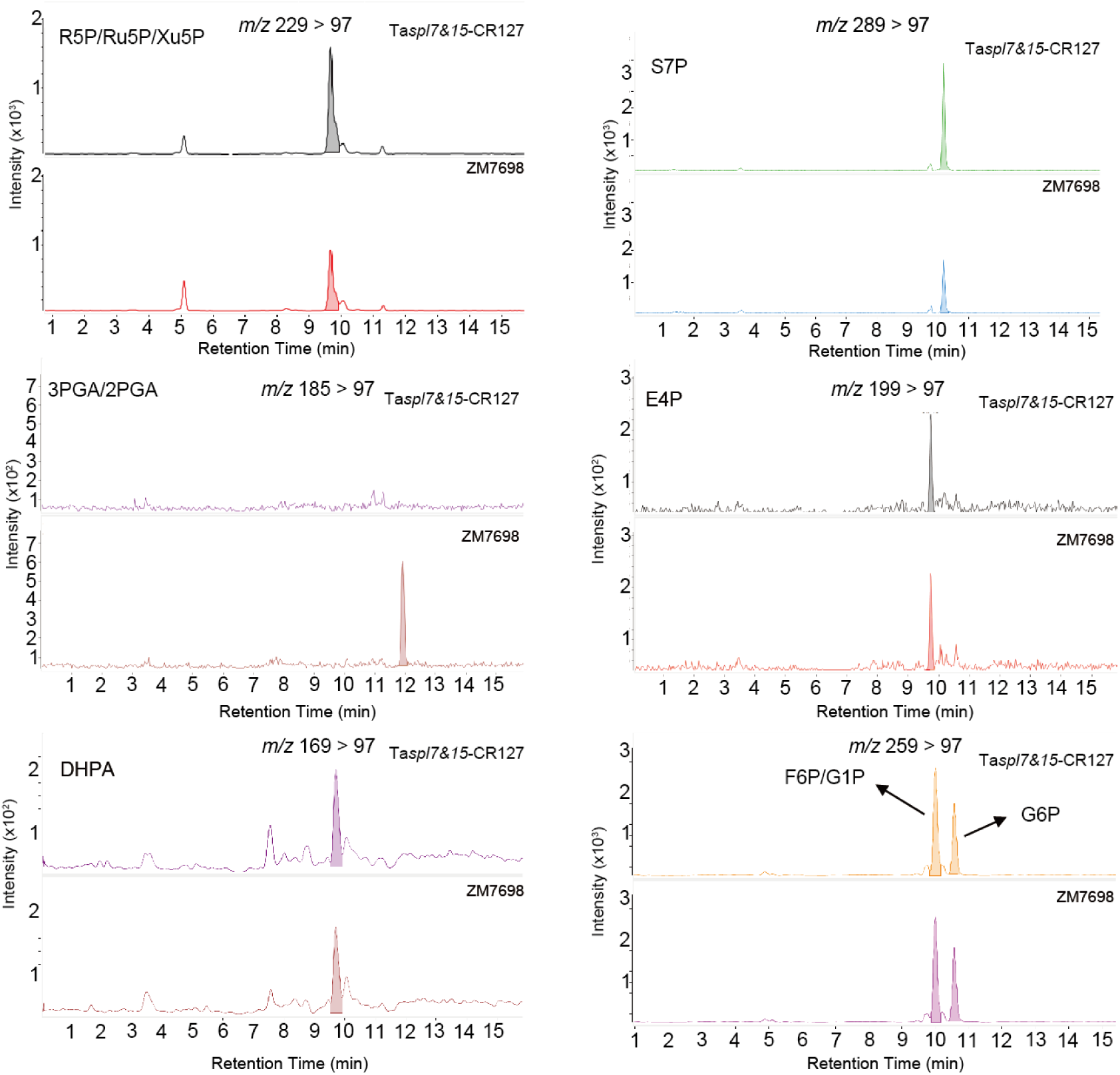
LC-QQQ-MS analysis of endogenous R5P/Ru5P/Xu5P, S7P, 3PGA/2PGA, E4P, DHPA, F6P/G1P and G6P from seedlings leaves of ZM7698 and *Taspl7&15*.

## Supplemental Tables

Table S1. Summary statistics of TAC-C libraries

Table S2. The intrachromosomal open chromatin loops

Table S3. The gene ID of C4 specific enzyme across four species

Table S4. The functional region used in this study

Table S5. Differentially expressed genes in *Taspl7&15* vs ZM7698

Table S6. Differentially expressed genes related to photosynthetic energy metabolism

Table S7. Chemicals and enzymes used in the TAC-C experiment

Table S8. Primers used in this study

